# Resolution enhancement with deblurring by pixel reassignment (DPR)

**DOI:** 10.1101/2023.07.24.550382

**Authors:** Bingying Zhao, Jerome Mertz

## Abstract

Improving the spatial resolution of a fluorescence microscope has been an ongoing challenge in the imaging community. To address this challenge, a variety of approaches have been taken, ranging from instrumentation development to image post-processing. An example of the latter is deconvolution, where images are numerically deblurred based on a knowledge of the microscope point spread function. However, deconvolution can easily lead to noise-amplification artifacts. Deblurring by post-processing can also lead to negativities or fail to conserve local linearity between sample and image. We describe here a simple image deblurring algorithm based on pixel reassignment that inherently avoids such artifacts and can be applied to general microscope modalities and fluorophore types. Our algorithm helps distinguish nearby fluorophores even when these are separated by distances smaller than the conventional resolution limit, helping facilitate, for example, the application of single-molecule localization microscopy in dense samples. We demonstrate the versatility and performance of our algorithm under a variety of imaging conditions.

## 1 Introduction

The spatial resolution of conventional fluorescence microscopy is limited to about half the emission wavelength because of diffraction.^1^ This limit can be surpassed using a variety of super-resolution approaches. For example, techniques such as STED,^2–6^ SIM,^7–13^ or ISM,^14–17^ generally require some form of scanning, either of a single excitation focus or of excitation patterns. Alternatively, scanning-free super-resolution can be achieved by exploiting the blinking nature of certain fluorophores, as popularized initially by PALM^18^ and STORM.^19^ In this latter approach, individual molecules are localized one by one based on the premise, typically, that the most likely location of the molecules is at the centroid of their respective emission point-spread-function (PSF). A distinct advantage of single molecule localization microscopy (SMLM) is that it can be implemented with a conventional camera-based fluorescence microscope, meaning that its barrier to entry is low and it can fully benefit from the most recent advances in camera technology (high quantum efficiency, low noise, massive pixel numbers, etc.). However, a key requirement of SMLM is that the imaged molecules are sparsely distributed, as ensured for example by photoactivation. This sparsity requirement implies, in turn, that several raw images, each sparse, are required to synthesize a final, less sparse, super-resolved image. Efforts have been made to partially alleviate this sparsity constraint in SMLM, such as SOFI,^20^ 3B analysis,^21^ DeconSTORM,^22^ and MUSICAL.^23^ Additional approaches not requiring molecular blinking have involved sparsifying the illumination, for example, with speckle illumination,^24^ or sparsifying the sample itself by tissue expansion ExM.^25, 26^ However, in all cases (excluding ExM) the requirement remains that several images, often thousands, are needed to produce an acceptable final image. As such, live-cell imaging is precluded and sparsity-based super-resolution approaches have been almost always limited to imaging fixed samples (though see Refs. 27–29).

In recent years, it has been noted that the sparsity constraint can be partially alleviated by pre-sharpening the raw images. Example algorithms are SRRF^30, 31^ and MSSR,^32^ which are freely available and easy to use. In contrast to DeconSTORM, these algorithms make only minimal assumptions about the emission PSF (radiality in the case of SRRF; convexity in the case of MSSR), and their application can substantially reduce the number of raw images required for SMLM. Indeed, when applied to denser images, only few images or even a single raw image can produce results quite comparable to much more time-consuming super-resolution approaches. However, these algorithms are not without drawbacks. SRRF and MSSR are both inherently highly nonlinear, meaning that additional steps are required to enforce a linear relation between sample and image brightness.^30–32^ Moreover, when applied to samples that are too dense, such as samples that exhibit features locally extended in 2D, those features tend to be hollowed out and only their edges or spines preserved, meaning that SRRF and MSSR are most applicable to samples containing only point- or line-like features smaller than the PSF, but provide poor fidelity otherwise.

We present an alternative image-sharpening approach that is similar to SRRF and MSSR, but has the advantage of inherently preserving image intensities and being more generally applicable. Like SRRF and MSSR, our approach can be applied to a wide variety of fluorescence microscopes where we make only minimal assumptions about the emission PSF, namely that the PSF centroid is located at its peak. Also, like SRRF and MSSR, our approach can be applied to a sequence of raw images, allowing a temporal analysis of blinking or fluctuation statistics, or it can be applied to only a few or even a single image. Our approach is based on post-processing by pixel reassignment, producing a deblurring effect similar to deconvolution but without some drawbacks associated with conventional deconvolution algorithms. We describe the basic principle of deblurring by pixel reassignment (DPR), and compare its performance to SRRF and MSSR both experimentally and with simulated data. Our DPR algorithm is made available as a Matlab function.

## 2 Principle of DPR

Fundamental to any linear imaging technique is the concept of a PSF: point sources in a sample produce light distributions at the imaging plane that are blurred by a convolution operation with the PSF. Because the width of the PSF is finite, so too is the image resolution. In principle, if the PSF is known exactly, the blurring caused by convolution can be undone numerically by deconvolution; however, in practice, such deblurring is hampered by fundamental limitations. For one, the Fourier transform of the PSF (or optical transfer function – OTF) provides a spatial frequency support inherently limited by the finite size of the microscope pupil, meaning that spatial frequencies beyond this diffraction limit are identically zero and cannot be recovered by deconvolution, even in principle (unless aided by assumptions^33–35^ such as sample analyticity, continuity, sparsity, etc.). Another limitation, no less fundamental, is the problem of noise. In conventional fluorescence microscopy, the OTF tapers to very small values as it approaches the diffraction limit, and falls below the shotnoise level generally well below this limit. As such, any attempt to amplify the high-frequency content of the OTF by deconvolution only ends up amplifying noise. This problem is particularly egregious with Wiener deconvolution,^36^ where noise amplification easily leads to unacceptable image mottling, and also with Richardson-Lucy (RL) deconvolution^37, 38^ when implemented with too many iterations. Regularization is required to dampen such noise-induced artifacts, often to the point that when applied to conventional fluorescence microscopy deconvolution only marginally improves resolution if at all (note that deconvolution fares better with non-conventional microscopies, such as SIM or ISM, where the OTF tapering near the diffraction limit is less severe).

The purpose of DPR is to perform PSF sharpening similar to deconvolution, but in a manner less prone to noise-induced artifacts and without the requirement of a full model for the PSF. Unlike Wiener deconvolution which is performed in Fourier space using a division operation, DPR operates entirely in real space with no division operation that can egregiously amplify noise. Unlike Richardson-Lucy deconvolution, DPR is non-iterative and can be performed in a single pass, without the need for an arbitrary iteration-termination criterion. DPR relies solely on pixel reassignment. As such, no negativities are possible in the final image reconstruction, as often encountered, for example, with Wiener deconvolution or image sharpening with a Laplacian filter.^39^ Moreover, intensity levels are rigorously conserved, with no requirement of additional procedures to ensure local linearity, as needed for example with SRRF, MSSR, or even SOFI.^40^

The basic principle of DPR is schematically depicted in Fig. 1 and described in more detail in Sec. 5. In brief, raw fluorescence images are first pre-conditioned by 1) performing global background subtraction, 2) normalizing to the overall maximum value in the image, and 3) re-mapping by interpolation to a coordinate system of grid period given by roughly 1/8th of the full width at half maximum (FWHM) of the PSF. The purpose of such pre-conditioning is to standardize the raw images prior to the application of DPR. The actual sharpening of the image is then performed by pixel reassignment, where intensities (pixel values) at each grid location (pixel) are reassigned to neighboring locations according to the direction and magnitude of the locally normalized image gradient (or, equivalently, the log-image gradient), scaled by a gain parameter. Because pixels are generally reassigned to off-grid locations, their pixel values are distributed to the nearest on-grid reassigned locations as weighted by their proximity (see Fig. S1 in the Supplementary Material). Finally, an assurance is included that pixels can be displaced no farther than 1.25 times the PSF FWHM width.

**Fig 1.**
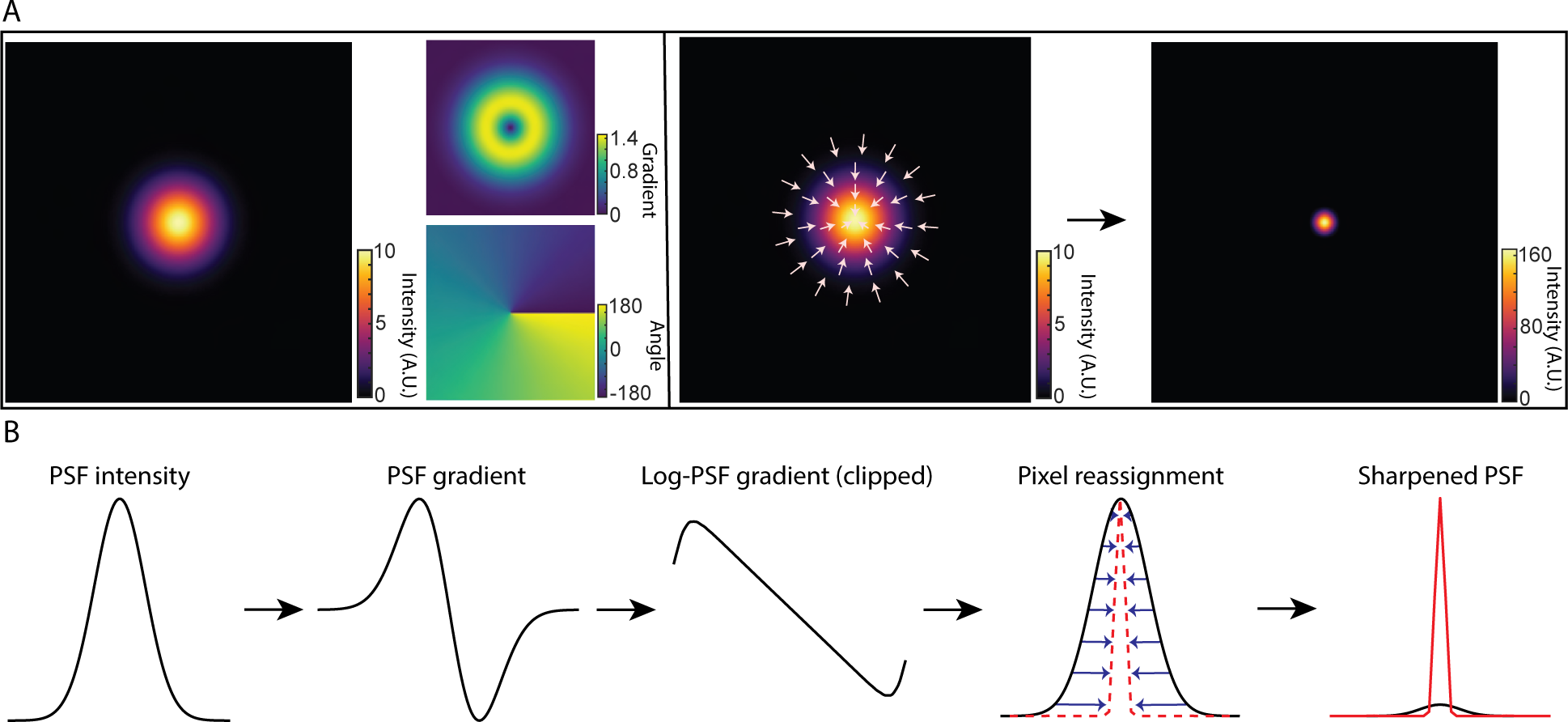
Principle of DPR. (a) From left to right: simulations of Gaussian PSF intensity and gradient maps (amplitude and direction); pixel reassignments; deblurred PSF image after application of DPR. (b) DPR workflow.

As a simple example, consider imaging a point source with a Gaussian PSF of Root-Mean-Square (RMS) width *σ*. The gradient of the log-PSF is then linear. That is, pixels are reassigned toward the PSF center exactly in proportion to their distance from the center, where the proportionality factor is selected by a gain parameter. The resultant sharpening of the PSF is substantial. For DPR gains 1 and 2, we find that the PSF widths are reduced by factors 4 and 7, respectively (see Fig. S2 in the Supplementary Material).

Conventionally, the resolution of a microscope is defined by its capacity to distinguish two point sources. More specifically, it is defined by the minimum separation distance required for two points to be resolved based on a predefined criterion, such as the Sparrow or Rayleigh criterion. We again consider the example of a Gaussian PSF, but now with two point sources. According to the Sparrow and Rayleigh criteria, the two points would have to be separated respectively by 2.2*σ* and 2.8*σ* to be resolvable. With the application of DPR, we find that this separation distance can be reduced. A clear dip between the two points by a factor of 0.74 is observed at a separation distance of 1.66*σ* for DPR gain 1, and an even smaller separation of 1.43*σ* for DPR gain 2. Indeed, we find that the two points remain resolvable down to separation distances of 1.36*σ* and 1.20*σ* for gains 1 and 2, respectively (Sparrow criterion), corresponding to resolution enhancements of 0.62 and 0.55 relative to the Sparrow limit, or, equivalently, 0.59 and 0.51 relative to the Rayleigh limit (see Fig. S3(b) in the Supplementary Material). It should be noted, however, that this enhanced capacity to resolve two nearby points is not entirely error-free. For example, when DPR is applied to points separated by less than about 1.9*σ*, the points begin to appear somewhat closer to each other than they actually are, with a relative error that increases with gain (see Fig. S3(c) in the Supplementary Material). That is, the choice of using DPR gain 1 or 2 (or other gain parameters) should be made at the user’s discretion bearing in mind this trade-off between resolution capacity and accuracy.

Similar resolution enhancement results are obtained when DPR is applied to two line objects. Here, we use raw data acquired by an Airyscan microscope obtained from^32, 41^ (Fig. 2(b)). In the raw image, lines separated by 150 nm cannot be resolved, whereas after the application of DPR with gains 1 and 2, they can be resolved at separations of 90 nm and 30 nm, respectively. The intensity profiles across the full set of line pairs for raw, DPR gain 1, and DPR gain 2 images are shown in Fig. S4(a) in the Supplementary Material. DPR images of the same sample acquired by conventional confocal microscopy^32, 41^ are shown in Fig. S4(b) in the Supplementary Material. In this case, lines separated by 210 nm cannot be resolved in the raw data set, whereas after application of DPR with gains 1 and 2, they can be resolved at separations of 120 nm and 90 nm, respectively.

**Fig 2.**
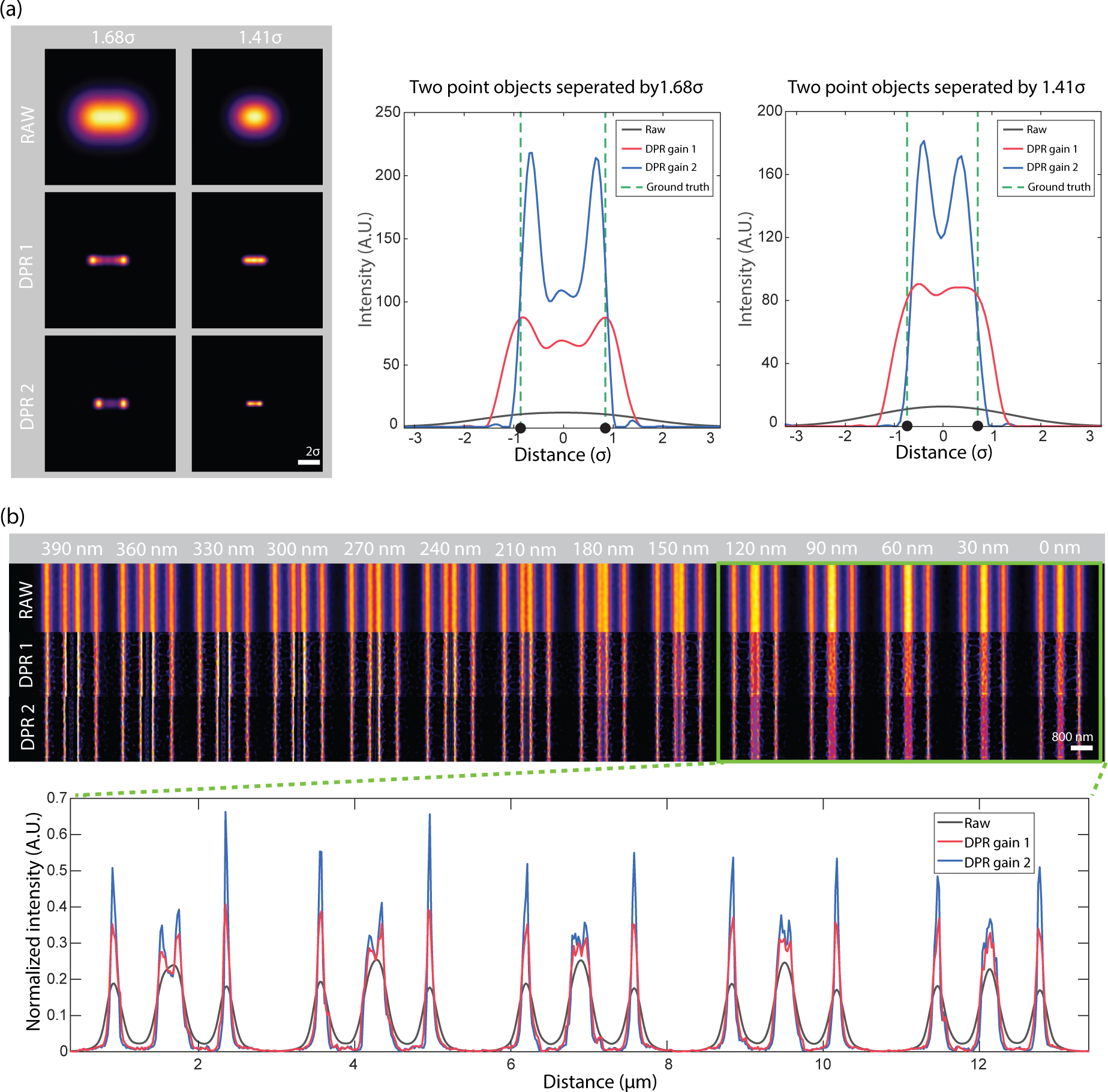
DPR resolution enhancement. (a) Simulation of DPR applied to two point objects separated by 1.68*σ* and 1.41*σ*. Left: images before (RAW) and after application of DPR gain 1 (DPR1) and DPR gain 2 (DPR2). Right: intensity plots across imaged points. Scale bar: 2*σ*. (b) DPR applied to fluorescent lines of increasing separation from 0 nm to 390 nm. Scale bar: 800 nm. PSF FWHM: 4 pixels. Local-minimum filter radius: 7 pixels.

To gain an appreciation of the effect of noise on DPR, we again simulated images of two point objects and two line objects, this time separated by 160 nm and imaged with a Gaussian PSF of RMS 84.93 nm. To these images we added shot noise (Poissonian) and additive camera readout noise (Gaussian) of different strengths, leading to SNR values of 5.0, 7.7, 14.1, and 20.3 (see Fig. S5 in the Supplementary Material). DPR gain 1 was applied to a stack of images, each with a different noise realization. The resulting resolution-enhanced images were then averaged over different numbers of frames (10, 20, and 40). Manifestly, the final image quality improves with increasing SNR and/or increasing numbers of frames averaged, as expected. Accordingly, the error in the measured separation between the two point objects and the two line objects as inferred by the separation between their peaks in the images also decreases (see Fig. S5(c) in the Supplementary Material). The images of line objects were less sensitive to noise, as evidenced by the relatively stable separation errors across various SNRs, but they exhibited somewhat higher separation error compared to the images of two point objects. These results are qualitative only. Nevertheless, they provide a rough indication of the increase in enhancement fidelity with SNR.

## 3 Results

### 3.1 DPR applied to single-molecule localization imaging

To demonstrate the resolution enhancement capacity of DPR with experimental data, we applied it to SMLM images. We used raw images made publicly available through the SMLM Challenge 2016,^42^ as these provide a convenient standardization benchmark. The experimental data consisted of a 4000-frame sequence of STORM images of microtubules labeled with Alexa567.

We applied DPR separately to each frame. Similar to SRRF and MSSR, we included in our DPR algorithm the possibility of temporal analysis of DPR-enhanced images. Here, the temporal analysis is simple, and consists either of averaging in time the DPR-enhanced images or calculating their variance in time (as is done, for example, with SOFI imaging of order 2). The results are shown in Figs. 3(a) and 3(b). As expected, DPR gain 2 leads to greater resolution enhancement than gain 1. Moreover, as expected, the temporal variance analysis leads to enhanced image contrast since it preferentially preserves fluctuating signals while removing non-fluctuating backgrounds. However, it should be noted that temporal variance analysis no longer preserves a linearity between sample and image strengths, as opposed to temporal averaging.

**Fig 3.**
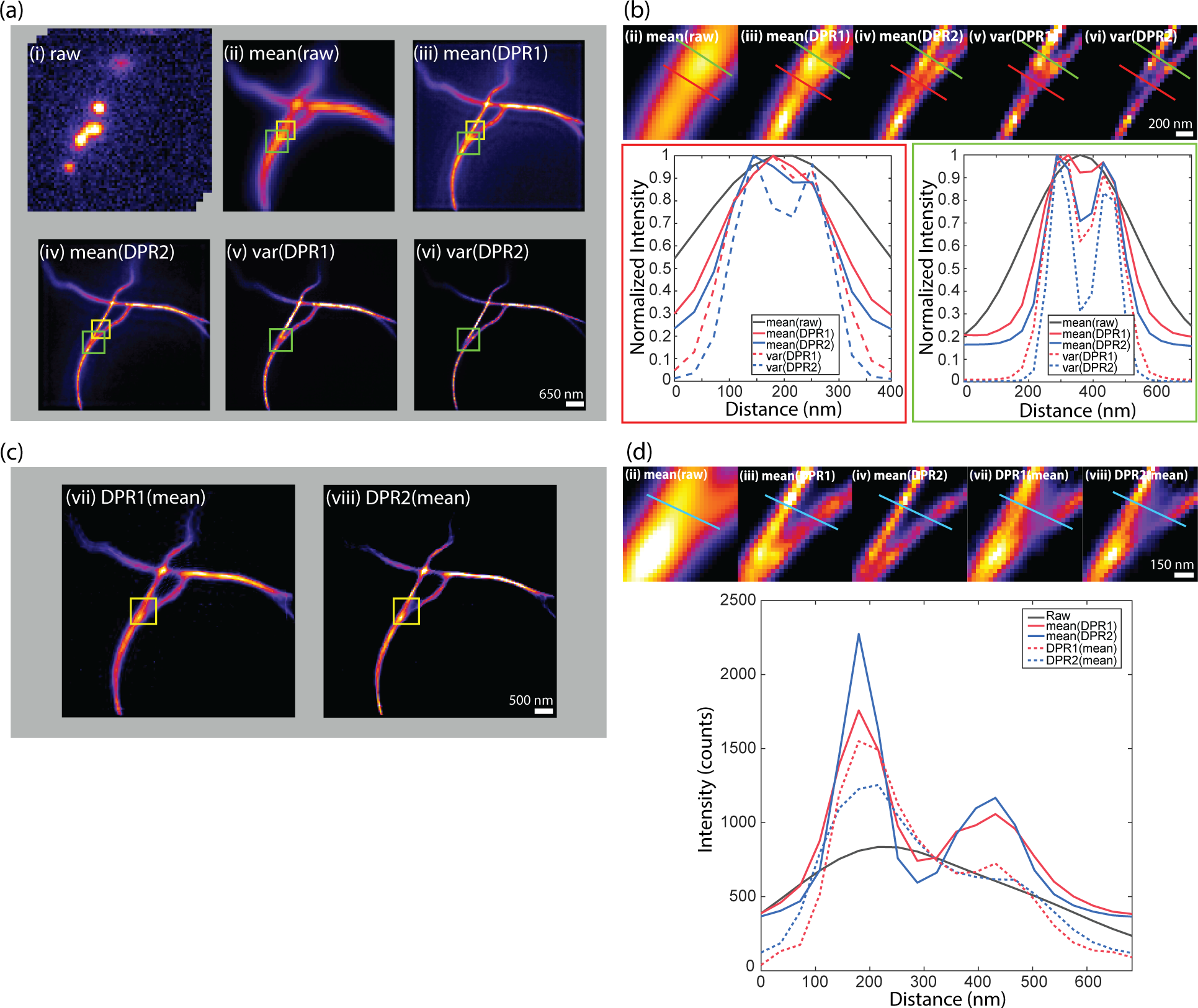
SMLM challenge 2016. (a) DPR applied to each frame in raw image stack, followed by temporal mean or variance. (i) raw image stack. (ii) mean of raw images. (iii) DPR gain 1 followed by mean. (iv) DPR gain 2 followed by mean. (v) DPR gain 1 followed variance. (vi) DPR gain 2 followed by variance. Scale bar: 650 nm. (b) Expanded regions of interest (ROIs) indicated by green square in (a). Bottom left: intensity distribution along red line in ROIs. Bottom right: intensity distribution along green line in ROIs. Scale bar: 200 nm. (c) Image mean followed by DPR, (vii) gain 1, (viii) gain 2. Scale bar: 500 nm. (d) Expanded ROIs indicated by yellow square in (c) and (a) (ii), (iii), and (iv) in (a). Right: intensity distribution along cyan line in ROIs. Scale bar: 150 nm. PSF FWHM: 2.7 pixels. Local-minimum filter radius: 5 pixels.

Interestingly, when a temporal average was applied to the raw images prior to the application of DPR (i.e. when the order of DPR and averaging was reversed – Figs. 3(c) and 3(d)), DPR continued to provide resolution enhancement, but not as effectively as when DPR was applied separately to each raw frame. The reason for this is clear. DPR relies on the presence of spatial structure in the image, which is largely washed out by averaging. In other words, similar to SRRF and MSSR, DPR is most effective when imaging sparse samples, as indeed is a requirement for SMLM.

### 3.2 DPR maintains imaging fidelity

DPR reassigns pixels according to their gradients. If the gradients are zero, the pixels remain in their initial position. That is, when imaging structures larger than the PSF that present gradients only around their edges but not within their interior, DPR sharpens only the edges while leaving the structure interiors unchanged. This differs, for example, from SRRF or MSSR, which erode or hollow out the interiors erroneously. DPR can thus be applied to more general imaging scenarios where samples contain both small and large structures. This is apparent, for example, when imaging a Siemens star target, as illustrated in Fig. S6 in the Supplementary Material, where neither SRRF nor MSSR accurately represents the widening of the star spokes. For example, when we applied NanoJ-SQUIRREL^43^ to the DPR-enhanced, MSSR-enhanced, and SRRF-enhanced Siemens star target images, we found resolution-scaled errors^43^ (RSEs) given by 53.5, 95.4, and 102.6 respectively; and resolution-scaled Pearson coefficients^43^ (RSPs) given by 0.92, 0.54, and 0.75 respectively. This improved fidelity is apparent, also, in other imaging scenarios. In Fig. 4, we show results obtained from the image of Alexa488-labeled bovine pulmonary artery endothelial (BPAE) cells (ThermoFisher, FluoCells) acquired with a conventional laser scanning confocal microscope (Olympus FLUOVIEW FV3000; objective: 40*×* air, 0.9NA; confocal pinhole set to 0.23x Airy units; PSF FWHM 256.4 nm; pixel size: 73.3 nm). While the intensity profiles along a single F-actin filament (red segments in Fig. 4) are sharpened roughly equally between SRRF, MSSR, and DPR, differences begin to appear for intensity profiles spanning nearby F-actin filaments (yellow segments in Fig. 4), or the imaging of larger structures (e.g. cluster in the blue box Fig. 4).

**Fig 4.**
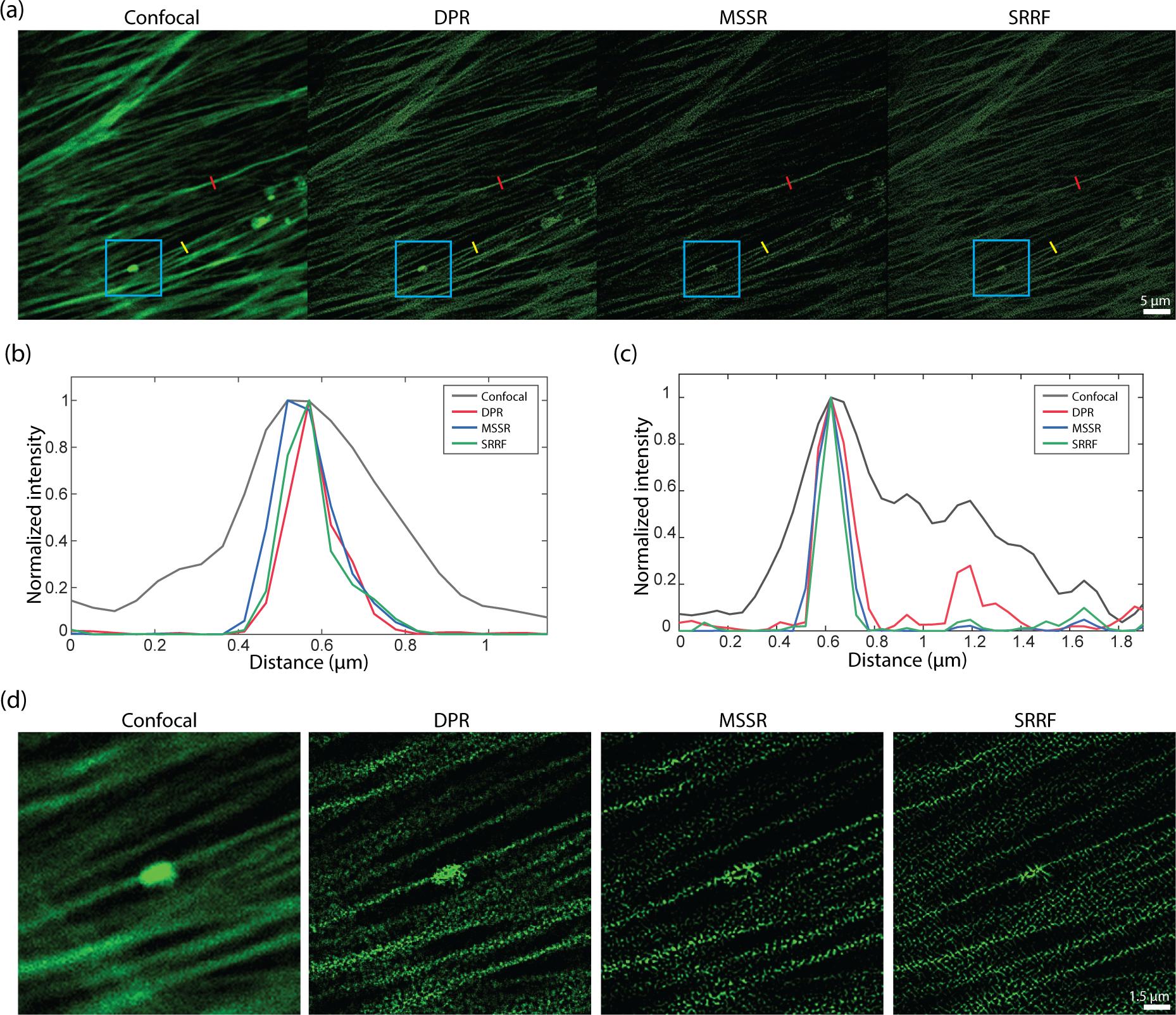
Comparison DPR, MSSR, and SRRF performances. (a) Images of BPAE cells acquired by a confocal microscope and after DPR gain 2, first order MSSR, and SRRF. DPR parameters: PSF FWHM 4 pixels, local-minimum filter radius 12 pixels, DPR gain 2. MSSR: PSF FWHM 4 pixels, magnification 2, order 1. SRRF: ring radius 0.5, magnification 2, axes 6. scale bar: 5 *µ*m. (b) Intensity profiles along single F-actin indicated by red lines in (c). (c) Intensity profiles along proximal F-actin filaments by yellow lines in (a). (d) Expanded ROIs indicated by cyan square in (a). scale bar: 1.5 *µ*m.

A difficulty when evaluating image fidelity is the need for a ground truth as a reference. To serve as a surrogate ground truth, we obtained images of BPAE cells with a state-of-the-art Nikon CS-WU1 spinning disk microscope equipped with a super-resolution module (SoRa) based on optical pixel reassignment^44^ (objective: 60*×* oil, 1.42NA; PSF FWHM 162.5 nm – conventional configuration, 114.9 nm - SoRa configuration; pixel size: 108.3 nm - conventional configuration, 27.1 nm - SoRa configuration), to which we additionally applied 20 iterations of RL deconvolution using the software supplied by the manufacturer. The same BPAE cells were also imaged at conventional (2*×* lower) resolution without the presence of the SoRa module. We then applied DPR, SRRF, and MSSR to the conventional resolution image for comparison (Fig. 5). As shown in the zoomed-in regions in Fig. 5(b), conventional confocal microscopy is not able to resolve two closely separated filaments, even after the application of RL-deconvolution. In contrast, DPR is easily able to resolve the filaments at both gains 1 and 2, providing images similar to the SoRa super-resolution ground-truth images. SRRF and MSSR also sharpened the filaments, but in the case of SRRF, the filaments remained difficult to resolve, while in both cases there was significant intensity dropout where the filaments disappeared altogether. When we applied NanoJ-SQUIRREL to compare the image fidelities of DPR, SRRF, and MSSR, we found the RSEs to be 11.35 for DPR gain 1, 11.12 for DPR gain 2, 19.36 for SRRF, and 18.88 for MSSR. Figure S7 in Supplementary Materials shows the RSE maps for DPR, SRRF, and MSSR. The RSPs were found to be 0.83 for DPR gain 1, 0.84 for DPR gain 2, 0.41 for SRRF, and 0.44 for MSSR. For these examples, DPR provides sharpened images with higher fidelity.

**Fig 5.**
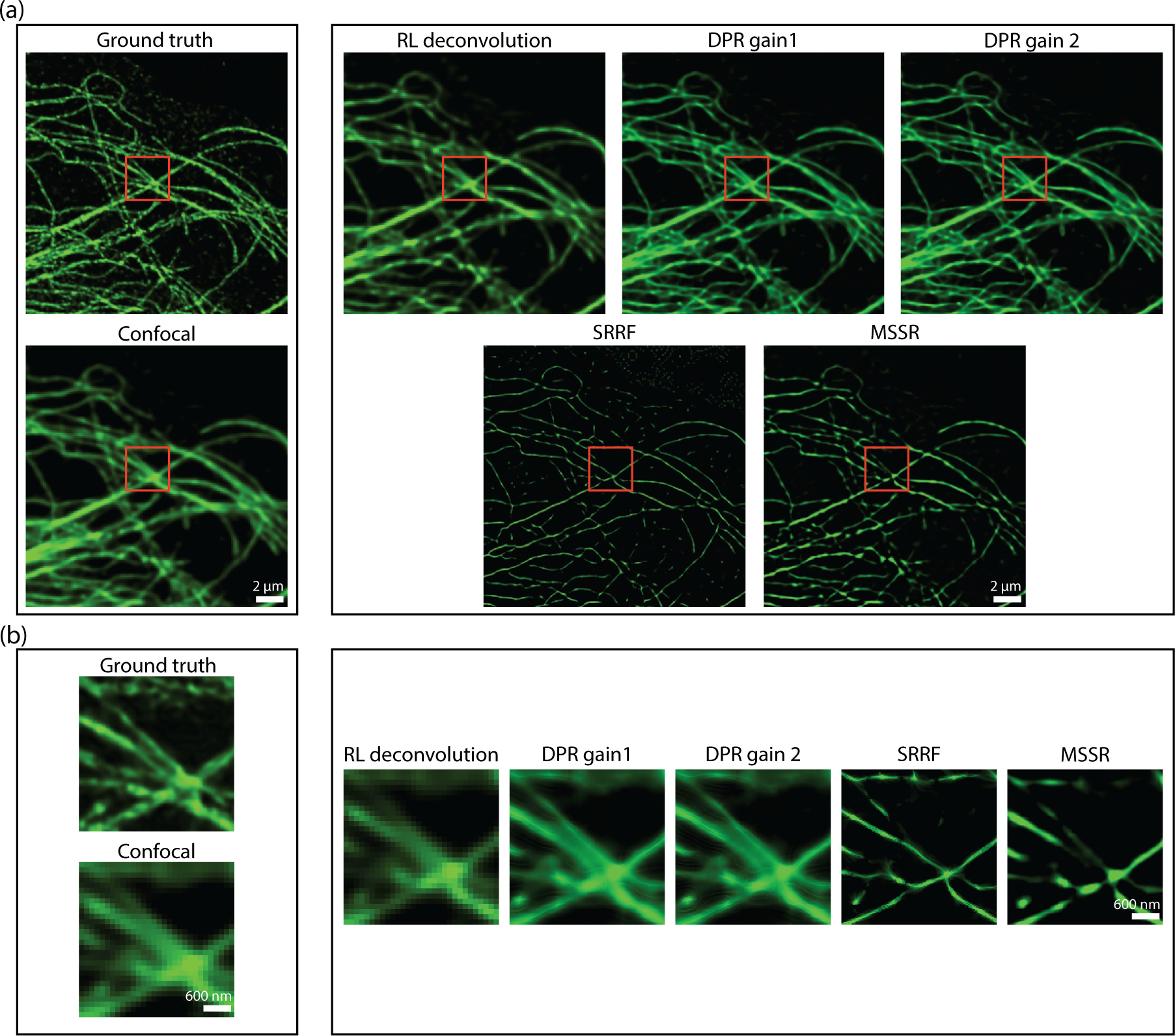
Comparison of DPR, SRRF, and MSSR with optical pixel reassignment and deconvolution. (a) Left: BPAE cells imaged using with of optical pixel reassignment and deconvolution with Nikon confocal microscope without (top) and with SoRa superresolution enhanced by RL deconvolution (20 iterations). Right: confocal image deblurred by RL deconvolution (20 iterations), DPR (gains 1 and 2), SRRF, and MSSR. DPR paramters: PSF FWHM 2 pixels, local-minimum filter radius: 40 pixels. MSSR parameters: PSF FWHM 2 pixels, magnification 4, order 1. SRRF: ring radius 0.5, magnification 4, axes 6. scale bar: 2 *µ*m. (b) The zoom-in ROIs indicated by red squares in (a). scale bar: 600 nm.

### 3.3 DPR applied to engineered cardiac tissue imaging

To demonstrate the ability of DPR to enhance image information, we performed structural imaging of engineered cardiac microbundles derived from human induced pluripotent stem cells (hiPSC), which have recently gained interest as model systems to study human cardiomyocytes (CMs).^45, 46^ We first imaged a monolayer of green fluorescent protein (GFP)-labeled hiPSC-CMs with a confocal microscope of sufficient resolution to reveal the z-discs of sarcomeres. This image serves as a ground truth reference. We then simulated a series of 45 conventional widefield images by numerically convolving the ground truth image with a low-resolution widefield PSF and adding simulated detection noise (shot noise and additive camera noise). DPR was applied to the conventional images, leading to the deblurred image shown in Fig. 6(a). Manifestly, this deblurred image much more closely resembles the ground truth image, as confirmed by the structural similarity index (SSIM):^47^ the SSIM between the conventional and ground truth images was 0.4, whereas, between the DPR and ground truth images, it was enhanced to 0.6. This increased fidelity is further validated by the pixel-wise error maps shown in Fig. 6(a), and by the line profiles through a sarcomere (cyan rectangle) showing that the application of DPR leads to better resolution of the z-discs, with the number and location of the z-discs being consistent with the ground truth image.

We also performed imaging of hiPSC cardiomyocyte tissue organoids (hiPSC-CMTs). Such imaging is more challenging because of the increased thickness of the organoids (about 400 *µ*m), which led to increased background and scattering-induced blurring, and also because of their irregular shapes which led to aberrations. Again we performed confocal imaging, this time at both low and high resolution (Figs. 6(b) and 6(c)), with lateral resolutions measured to be 1.5 *µ*m and 0.5 *µ*m, respectively, based on the FWHM of sub-diffraction-sized fluorescent beads. As expected, low-resolution imaging failed to clearly resolve the z-discs (the separation between z-discs is in the range of 1.8-2.0 *µ*m, depending on sarcomere maturity). However, when DPR was applied to the low-resolution image the z-discs became resolvable (Fig. 6(b)). DPR was further applied to the high-resolution image, resulting in an even greater enhancement of resolution (Fig. 6(c)).

**Fig 6.**
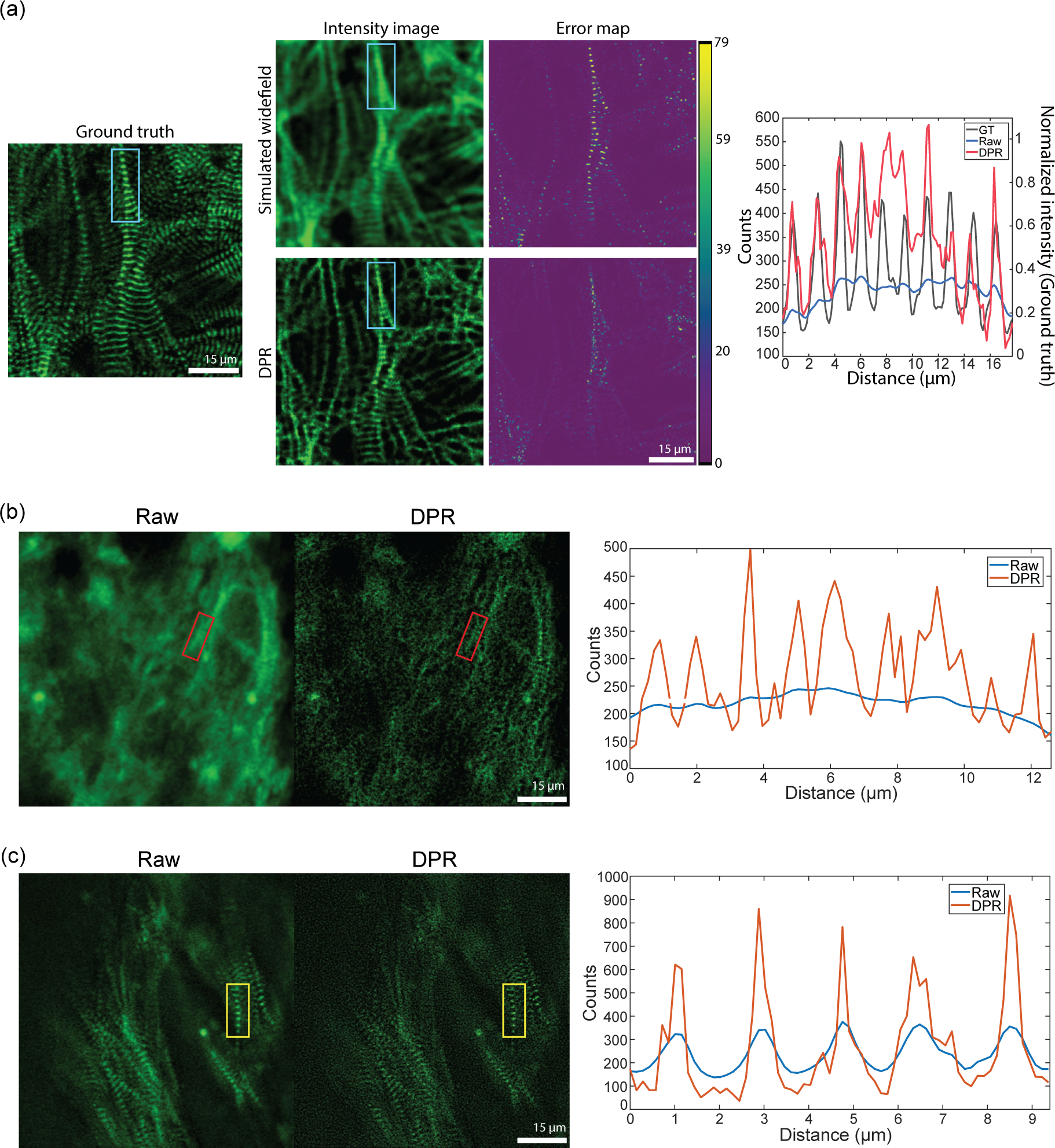
Engineered cardiac tissue imaging. (a) DPR gain 1 applied to simulated ground-truth widefield images of mono-layer hiPSC-CMs derived from experimental images acquired by a confocal microscope. Left: simulated ground truth. Middle: simulated widefield intensity image without (top) and with (bottom) application of DPR, and corresponding error maps. Right: intensity profile along sarcomere chain indicated by the cyan rectangle. PSF FWHM: 4 pixels, local-minimum filter radius: 7 pixels. (b) DPR gain 1 applied to experimental low-resolution images of hiPSC-CMT organoids. Left: confocal image. Middle: DPR-enhanced image. Right: intensity profile of sarcomere chain indicated by the red rectangle. PSF FWHM: 4 pixels, local-minimum filter radius: 7 pixels. (c) DPR gain 1 applied to experimental high-resolution images of hiPSC-CMT. Left: confocal image. Middle: DPR-enhanced image. Right: intensity profile of the sarcomere chain indicated by the yellow rectangle. PSF FWHM: 2 pixels, local-minimum filter radius: 4 pixels. Scale bar: 15 *µ*m.

### 3.4 DPR applied to volumetric zebrafish imaging

In recent years, there has been a push to develop microscopes capable of imaging populations of cells within extended volumes at high spatiotemporal resolution. One such microscope is based on confocal imaging with multiple axially distributed pinholes, enabling simultaneous multiplane imaging.^48, 49^ However, in its most simple implementation, multi-z confocal microscopy is based on low-NA illumination and provides only limited spatial resolution, roughly 2.6 *µ*m lateral and 15 *µ*m axial. While such resolution is adequate for distinguishing neuronal somata in animals like mice and zebrafish, it is inadequate for distinguishing, for example, nearby dendritic processes. To demonstrate the applicability of DPR to different types of microscopes, we applied it here to zebrafish images acquired with a multi-z confocal microscope essentially identical to that described in Ref. 48 (objective: 16X water, 0.8NA; pixel size: 0.5 *µ*m.). We imaged neuronal process in the head and tail regions of a transgenic zebrafish larva expressing GFP at 9 days post fertilization (dpf). Both regions were imaged with four planes separated by 20 *µ*m, within a volume of 256*×*256*×*60 *µ*m^3^. Without DPR, the axons in the brain region (Fig. 7(a)) are difficult to resolve, as expected, since this region is densely labeled; whereas in the tail region (Fig. 7(b)) where axons are more sparsely distributed and there is less background, the axons are resolvable but blurred. When DPR is applied, the axons in both the brain and tail regions become deblurred and clearly resolvable, enabling a cleaner separation between image planes (Figs. 7(a) and 7(d)). We note that these results are more qualitative than quantitative since no ground truth was available for reference. Nevertheless, we did compare our DPR results with raw images obtained by a different multi-z system equipped with a diffractive optical element in the excitation path to enable higher-resolution imaging (0.50 *µ*m lateral and 3.6 *µ*m axial).^50^ A comparison is shown in Fig. 7(d) (different fish, different tail regions – see also Fig. S9 in the Supplementary Material), illustrating the qualitative similarity between low-resolution multi-z images enhanced with DPR and directly-acquired higher resolution images.

**Fig 7.**
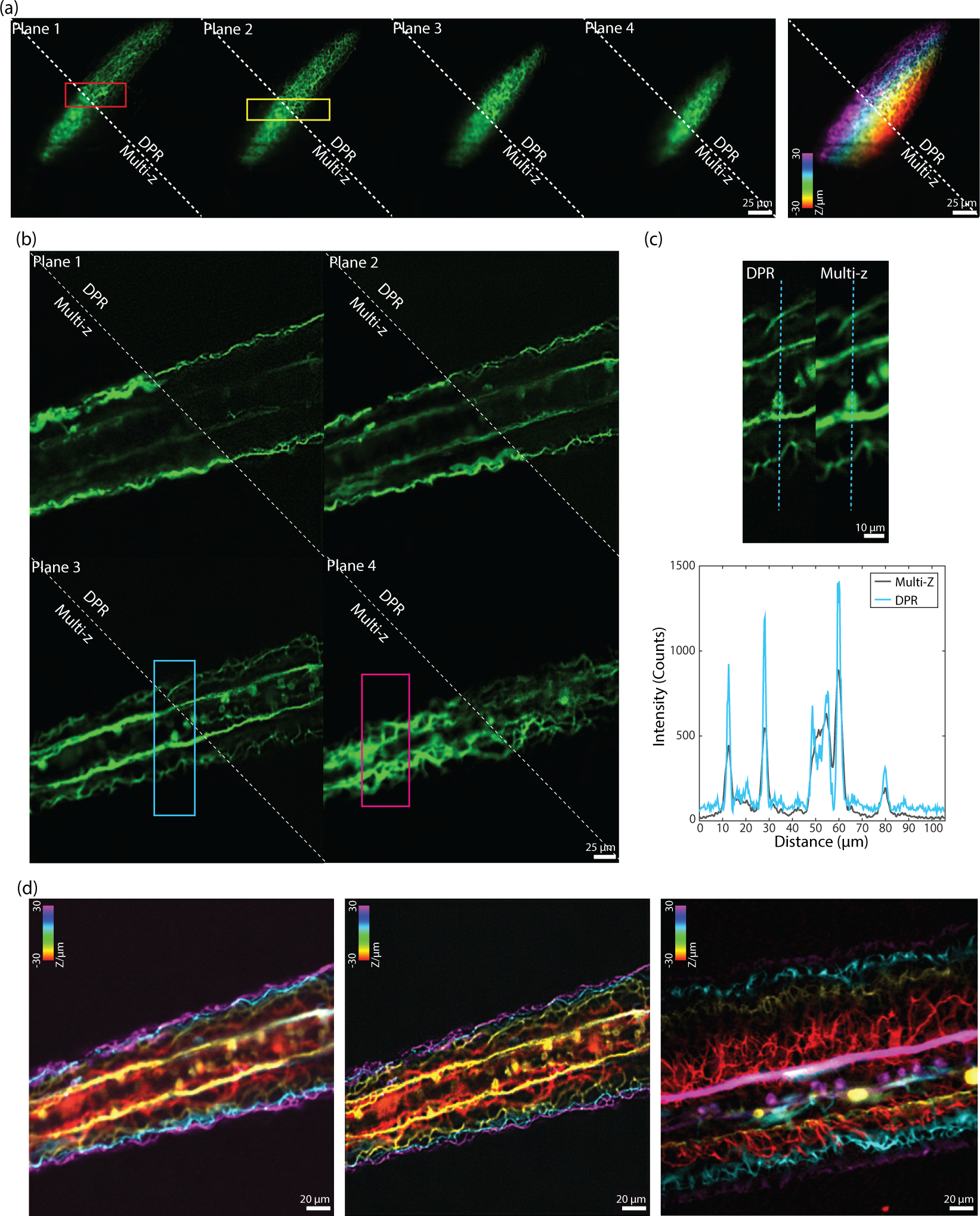
Multi-z confocal zebrafish imaging. (a) *In-vivo* raw and DPR-enhanced (gain 1) multiplane images of the brain region in a zebrafish 9 dpf. Left: four image planes. Right: merged with colors corresponding to depth. ROI indicated by the red and yellow rectangles in (a) are shown in Fig. S8 in the Supplementary Material. Scale bar: 25 *µ*m. (b) Raw and DPR-enhanced images multiplane images of the zebrafish tail region. Scale bar: 25 *µ*m. (c) ROI indicated by the cyan rectangle in (b), and intensity profile along the cyan dashed line. ROI indicated by the magenta rectangles in (b) are shown in Fig. S8 in the Supplementary Material. Scale bar: 10 *µ*m. (d) Merged multiplane images in the tail region of a zebrafish. Left: raw low-resolution multiplane images. Middle: DPR-enhanced low-resolution multiplane images. Right: raw high-resolution multiplane images (different fish and tail regions). PSF FWHM: 5, local-minimum filter radius: 25 pixels. Scale bar: 20 *µ*m. Plane 1-4: the deepest to the shallowest. Inter-plane separation: 20 *µ*m.

## 4 Discussion

The purpose of DPR is to help counteract the blurring induced by the PSF of a fluorescence microscope. The underlying assumption of DPR is that fluorescent sources are located by their associated PSF centroids, which are found by hill-climbing directed by local intensity gradients.^51, 52^ When applied to individual fluorescence images, DPR helps distinguish nearby sources even when these are separated by distances smaller than the Sparrow or Rayleigh limit. In other words, DPR can provide resolution enhancement even in densely labeled samples. Such resolution enhancement is akin to image sharpening, with the advantage that DPR is performed in real space rather than Fourier space, and that local intensities are inherently preserved and negativities are impossible (Table. S1 in the Supplementary Materials).

To define what is meant by the term *local* here, we can directly compare the intensities of raw and DPR-enhanced images. When both images are spatially filtered by average blurring, the differences between their intensities become increasingly small with increasing kernel size of the average filter (see Fig. S10). Indeed, the relative differences, as characterized by the difference standard deviations, drop to less than 7.7% when the kernel size is smaller than about 4.5 times the PSF FWHM (corresponding to about 36 sub-pixels). In other words, on scales larger than 4.5 PSF widths, the raw and DPR images are essentially identical. It is only on scales smaller than 4.5 PSF widths that deviations between the two images begin to appear owing to the image sharpening induced by DPR. The image sharpening, which is inherently local, thus preserves intensities on this scale. If, in addition, the sample can be regarded as sparse, either by assumption or because of fluorophore intensity fluctuation statistics (imposed or passive), the enhanced capacity of DPR to distinguish fluorophores can help reduce the number of images required for SMLM-type superresolution.

Of course, no deblurring strategy is immune to noise, and the same is true for DPR. However, DPR presents the advantage that noise cannot be amplified as it can be, for example, with Wiener or RL deconvolution, both of which require some form of noise regularization (in the case of RL, iteration termination is equivalent to regularization). DPR requires neither regularization nor even an exact model of the PSF. As such, DPR resembles SRRF and MSSR, but with the advantage of simpler implementation and more general applicability to samples possessing extended features.

Finally, our DPR algorithm is made available here as a MATLAB function compatible with either Windows or macOS. An example time to process a stack of 128*×*128*×*100 images obtained with a microscope whose PSF FWHM is 2 pixels (upscaled to 533*×*533*×*100) when run on an Intel i7-9800X computer equipped with an NVIDIA GeForce GTX 970 GPU are 2.32 s. A similar run time is found when using a MacBook Pro with Apple M1 Pro (2.08 s with MATLAB).

Because of its ease of use, speed, and versatility, we believe DPR can be of general utility to the bio-imaging community.

## 5 Methods

### 5.1 DPR algorithm

Figure S9 in the Supplementary Material illustrates the overall workflow of DPR, and Algorithm S1 provides more details in the form of a pseudo-code. To begin, we subtract the global minimum from each raw image to remove uniform background and camera offset (if any). These background-subtracted images *I_in_*serve as the inputs to our algorithm.

Next, we must establish a vector map to guide the pixel reassignment process. This is done in steps. First, we perform local equalization of *I_in_* by applying a local-minimum filter and subtracting it from *I_in_*, obtaining *I_eq_*. The radius of the local-minimum filter is user-defined. Typically, we use a radius 10*×* the Airy disk (default), though this can be made bigger to preserve larger sample structures. Next, both *I_in_* and *I_eq_* are mapped onto a grid of period equal to 8*×* the estimated FWHM of the PSF. This mapping is performed by spline interpolation.^53^ The images are zero-padded to prevent errors due to out-of-image pixel reassignments, where the zero-padding is removed at the end of DPR. We also divide *I_eq_* by a lowpass-filtered version of itself, so as to locally normalize *I_eq_*. It is from this locally-normalized *I_eq_* that the pixel-reassignment vector map is derived.

The reassignment vector map for DPR is obtained by first calculating the gradients of *I_eq_* along the x and y directions using conventional 3 *×* 3 Sobel gradient filters.^54^ The resulting gradient vectors for each pixel are then normalized to the pixel values themselves (equivalent to a log-image gradient), and multiplied by a scalar gain. The comparison of using image gradients and log-image gradients for pixel reassignment is illustrated in Fig. S11 in the Supplementary Material. The gain is user-defined. For our results, we used gains of 1 or 2. Note that the length of the reassignment vector is limited in part by a small offset applied to the equalized image that ensures the normalized image gradients remain finite. In addition, a hard limit of 10 pixels (1.25 times the PSF FWHM) is imposed: reassignment vectors of lengths longer than 10 pixels are ignored, and the pixels are left unchanged.

Pixel reassignment consists of numerically displacing the intensity values in the input image from their initial grid position to new reassigned positions according to their associated pixel reassignment vector. In general, as shown in Fig. S1 in the Supplementary Material, the new reassigned positions are off-grid. The pixel values are then partitioned to the nearest four on-grid locations as weighted by their proximity to these locations, as described in more detail in Algorithm S1 of the Supplementary Material.

Reassignment is performed pixel-by-pixel across the entire input image, leading to a final output DPR image. In the event that a time sequence of the image is processed, the output DPR sequence can be temporally analyzed (for example, by calculating the temporal average or variance) if desired. Note that an input parameter for our DPR algorithm is the estimated PSF size. When using the Olympus FV3000 microscope, we obtained this from the manufacturer’s software. When using our home-built confocal microscopes, we measured this with 200 nm fluorescent beads (Phosphorex). When using the SoRa microscope, we used the estimated PSF for confocal microscopy. The PSF size needs not be exact, and may be estimated from the conventional Rayleigh resolution limit given by^55^

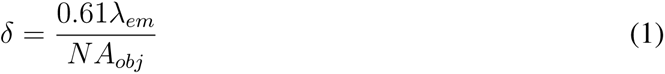

where *λ_em_*is the emission wavelength and NA*_obj_* is the numerical aperture of the objective.

### 5.2 Simulated data

Simulated widefield images of two point objects and two line objects separated by 160 nm were used to evaluate the separation accuracy of DPR using Gaussian PSF of standard deviation 84.93 nm. The images were rendered on a 40 nm grid. Poisson and Gaussian noise were added to simulate different SNRs, with SNR being calculated by

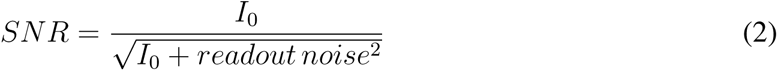

A temporal stack of 45 image frames with independent noises was generated for each SNR.

Simulated widefield images of the sarcomere ground truth were produced based on a Gaussian PSF model of standard deviation 0.85 *µ*m. Poisson noise and Gaussian noise were added to the images. A temporal stack of 45 image frames of images was then generated.

### 5.3 Engineered heart tissue preparation

hiPSCs from the PGP1 parent line (derived from PGP1 donor from the Personal Genome Project) with an endogenous GFP tag on the sarcomere gene TTN^56^ were maintained in mTeSR1 (Stem-Cell) on Matrigel (Fisher) mixed 1:100 in DMEM/F-12 (Fisher) and split using Accutase (Fisher) at 60–90% confluence. hiPSCs were differentiated into monolayer hiPSC-CMs by the Wnt signaling pathway.^57^ Once cells were beating, hiPSC-CMs were purified using RPMI no-glucose media (Fisher) with 4 mM sodium DL lactate solution (Sigma) for 2–5 days. Following selection, cells were replated and maintained in RPMI with 1:50 B-27 Supplement (Fisher) on 10 *µ*g/mL fibronectin (Fisher)-coated plates until day 30.

3D hiPSC-CMTs devices with tissue wells, each containing two cylindrical micropillars with spherical caps, were cast in PDMS from a 3D-printed mold (Protolabs).^58^ A total of 60,000 cells per CMT, 90% hiPSC-CMs and 10% Normal Human Ventricular Cardiac Fibroblasts (NHCF-V) were mixed in 7.5 µL of an ECM solution, 4 mg/mL human fibrinogen (Sigma), 10% Matrigel (Corning), 1.6 mg/mL thrombin (Sigma), 5 *µ*M Y-27632 (Tocris), and 33 *µ*g/mL aprotinin (Sigma). The cell-ECM mixture was pipetted into each well, and after polymerization for 5 min, growth media containing high-glucose DMEM (Fisher) supplemented with 10% fetal bovine serum (Sigma), 1% penicillin–streptomycin (Fisher), 1% nonessential amino acids (Fisher), 1% GlutaMAX (Fisher), 5 *µ*M Y-27632, and 33 *µ*g/mL aprotinin was added and replaced every other day. Y-27632 was removed 2 days after seeding, and aprotinin was decreased to 16 *µ*g/mL after 7 days. Next, CMTs were fixed in 4% PFA (Fisher) for 30 min, washed 3x with PBS, and stored in PBS at 4*^◦^*C.

### 5.4 Zebrafish preparation

All procedures were approved by the Institutional Animal Care and Use Committee (IACUC) at Boston University, and practices were consistent with the Guide for the Care and Use of Laboratory Animals and the Animal Welfare Act. For the *in-vivo* structural imaging of zebrafish, transgenic zebrafish embryos (isl2b:Gal4 UAS:Dendra) expressing GFP were maintained in filtered water from an aquarium at 28.5*^◦^*C on a 14 - 10 hr light-dark cycle. Zebrafish larvae at 9 days post-fertilization (dpf) were used for imaging. The larvae were embedded in 5% low-melting-point agarose (Sigma) in a 55 mm petri dish. After agarose solidification, the petri dish was filled with filtered water from the aquarium.

### 5.5 hiPSC-CMTs imaging

The hiPSC-CMTs were imaged with a custom confocal microscope, essentially identical to that described in Ref. 48, but with adjustable illumination NA (0.2NA with 300 *µ*m confocal pinhole for low-resolution imaging and 0.8NA with 150 *µ*m confocal pinhole for high-resolution imaging) and single-plane detection. The measured PSF FWHMs were 1.5 *µ*m for low-resolution imaging and 0.5 *µ*m for high-resolution using the 16*×* objective (Nikon CFI LWD Plan Fluorite Water 16*×*, 0.8NA). Pixel sizes: 0.3 *µ*m.

### 5.6 DPR, MSSR, and SRRF parameters

The parameters used for DPR, MSSR, and SRRF for our results can be found in Table S2 in the Supplementary Materials.

### 5.7 Error map calculation

Error map calculation is realized by a custom script written in Matlab R2021b. Pixelated differences between the images and the ground truth are directly measured by subtraction and saved as an error map.

### 5.8 SSIM calculation

The SSIM calculation is realized by the SSIM function in Matlab R2021b. Exponents for luminance, contrast, and structural terms are set as [1, 1, 1] (default values). Standard deviation of the isotropic Gaussian function is set as 1.5 (default value).

## Code, Data, and Materials Availability

Data underlying the results presented in this paper are not publicly available at this time but may be obtained from the authors upon reasonable request.

The DPR Matlab package is available on Github (https://github.com/biomicroscopy/DPR-Project).

## Acknowledgments

We thank Ichun Anderson Chen and Shuqi Zheng for assistance in the confocal imaging of BPAE cells and Zahid Yaqoob for assistance in the SoRa microscope and helpful discussion. We also thank Florian Engert and Paula Pflitsch for supplying the zebrafish (isl2b:Gal4 UAS: Dendra) and Joshua Lee for help in the hiPSC-CMTs sample preparation. This work was funded by National Science Foundation (EEC-1647837, 2215990) and National Institutes of Health (R01EB029171, R01NS116139).

## References

1 J. Mertz, Introduction to optical microscopy, Cambridge University Press (2019).

2 J. Wichmann and S. W. Hell, “Breaking the diffraction resolution limit by stimulated emission: stimulated-emission-depletion fluorescence microscopy,” Optics Letters, Vol. 19, Issue 11, pp. 780–782 19, 780–782 (1994).

3 T. A. Klar, S. Jakobs, M. Dyba, et al., “Fluorescence microscopy with diffraction resolution barrier broken by stimulated emission,” Proceedings of the National Academy of Sciences of the United States of America 97, 8206–8210 (2000).

4 S. W. Hell, “Toward fluorescence nanoscopy,” Nature Biotechnology 2003 21:11 21, 1347–1355 (2003).

5 K. I. Willig, S. O. Rizzoli, V. Westphal, et al., “Sted microscopy reveals that synaptotagmin remains clustered after synaptic vesicle exocytosis,” Nature 440, 935–939 (2006).

6 E. Rittweger, K. Y. Han, S. E. Irvine, et al., “Sted microscopy reveals crystal colour centres with nanometric resolution,” Nature Photonics 3, 144–147 (2009).

7 M. G. Gustafsson, “Surpassing the lateral resolution limit by a factor of two using structured illumination microscopy,” Journal of Microscopy 198 (2000).

8 R. Heintzmann, T. M. Jovin, and C. Cremer, “Saturated patterned excitation microscopy—a concept for optical resolution improvement,” JOSA A, Vol. 19, Issue 8, pp. 1599–1609 19, 1599–1609 (2002).

9 M. G. L. Gustafsson, “Nonlinear structured-illumination microscopy: Wide-field fluorescence imaging with theoretically unlimited resolution,” Proceedings of the National Academy of Sciences 102(37), 13081–13086 (2005).

10 P. Kner, B. B. Chhun, E. R. Griffis, et al., “Super-resolution video microscopy of live cells by structured illumination,” Nature methods 6(5), 339–342 (2009).

11 M. G. Gustafsson, L. Shao, P. M. Carlton, et al., “Three-dimensional resolution doubling in wide-field fluorescence microscopy by structured illumination,” Biophysical Journal 94, 4957–4970 (2008).

12 L. Schermelleh, P. M. Carlton, S. Haase, et al., “Subdiffraction multicolor imaging of the nuclear periphery with 3d structured illumination microscopy,” Science 320(5881), 1332–1336 (2008).

13 D. Li, L. Shao, B. C. Chen, et al., “Extended-resolution structured illumination imaging of endocytic and cytoskeletal dynamics,” Science 349 (2015).

14 C. B. Müller and J. Enderlein, “Image scanning microscopy,” Physical review letters 104(19), 198101 (2010).

15 S. Roth, C. J. Sheppard, K. Wicker, et al., “Optical photon reassignment microscopy (opra),” Optical Nanoscopy 2, 1–6 (2013).

16 G. M. De Luca, R. M. Breedijk, R. A. Brandt, et al., “Re-scan confocal microscopy: scanning twice for better resolution,” Biomedical optics express 4(11), 2644–2656 (2013).

17 C. J. Sheppard, S. B. Mehta, and R. Heintzmann, “Superresolution by image scanning microscopy using pixel reassignment,” Optics letters 38(15), 2889–2892 (2013).

18 E. Betzig, G. H. Patterson, R. Sougrat, et al., “Imaging intracellular fluorescent proteins at nanometer resolution,” Science 313, 1642–1645 (2006).

19 M. J. Rust, M. Bates, and X. Zhuang, “Sub-diffraction-limit imaging by stochastic optical reconstruction microscopy (storm),” Nature Methods 2006 3:10 3, 793–796 (2006).

20 T. Dertinger, R. Colyera, G. Iyer, et al., “Fast, background-free, 3d super-resolution optical fluctuation imaging (sofi),” Proceedings of the National Academy of Sciences of the United States of America 106, 22287–22292 (2009).

21 S. Cox, E. Rosten, J. Monypenny, et al., “Bayesian localization microscopy reveals nanoscale podosome dynamics,” Nature Methods 2011 9:2 9, 195–200 (2011).

22 E. A. Mukamel, H. Babcock, and X. Zhuang, “Statistical deconvolution for superresolution fluorescence microscopy,” Biophysical Journal 102, 2391–2400 (2012).

23 K. Agarwal and R. Machá, “Multiple signal classification algorithm for super-resolution fluorescence microscopy,” Nature Communications 2016 7:1 7, 1–9 (2016).

24 T. Mangeat, S. Labouesse, M. Allain, et al., “Super-resolved live-cell imaging using random illumination microscopy,” Cell Reports Methods 1(1), 100009 (2021).

25 F. Chen, P. W. Tillberg, and E. S. Boyden, “Expansion microscopy,” Science 347, 543–548 (2015).

26 I. Cho, J. Y. Seo, and J. Chang, “Expansion microscopy,” Journal of Microscopy 271, 123–128 (2018).

27 S. A. Jones, S. H. Shim, J. He, et al., “Fast, three-dimensional super-resolution imaging of live cells,” Nature Methods 2011 8:6 8, 499–505 (2011).

28 S. van de Linde, M. Heilemann, and M. Sauer, “Live-cell super-resolution imaging with synthetic fluorophores,” Annual review of physical chemistry 63, 519–540 (2012).

29 H. Takakura, Y. Zhang, R. S. Erdmann, et al., “Long time-lapse nanoscopy with spontaneously blinking membrane probes,” Nature Biotechnology 2017 35:8 35, 773–780 (2017).

30 N. Gustafsson, S. Culley, G. Ashdown, et al., “Fast live-cell conventional fluorophore nanoscopy with imagej through super-resolution radial fluctuations,” Nature communications 7(1), 1–9 (2016).

31 R. F. Laine, H. S. Heil, S. Coelho, et al., “High-fidelity 3d live-cell nanoscopy through data-driven enhanced super-resolution radial fluctuation,” bioRxiv (2022).

32 E. Torres-García, R. Pinto-Cámara, A. Linares, et al., “Extending resolution within a single imaging frame,” Nature Communications 13(1), 7452 (2022).

33 S. Hugelier, J. J. De Rooi, R. Bernex, et al., “Sparse deconvolution of high-density super-resolution images,” Scientific reports 6(1), 21413 (2016).

34 X. Huang, J. Fan, L. Li, et al., “Fast, long-term, super-resolution imaging with hessian structured illumination microscopy,” Nature biotechnology 36(5), 451–459 (2018).

35 W. Zhao, S. Zhao, L. Li, et al., “Sparse deconvolution improves the resolution of live-cell super-resolution fluorescence microscopy,” Nature biotechnology 40(4), 606–617 (2022).

36 N. Wiener, Extrapolation, interpolation, and smoothing of stationary time series: with engineering applications, MIT press Cambridge, MA (1949).

37 W. H. Richardson, “Bayesian-based iterative method of image restoration*∗*,” J. Opt. Soc. Am. 62, 55–59 (1972).

38 L. B. Lucy, “An iterative technique for the rectification of observed distributions,” The astronomical journal 79, 745 (1974).

39 M. Aubry, S. Paris, S. W. Hasinoff, et al., “Fast local laplacian filters: Theory and applications,” ACM Trans. Graph. 33 (2014).

40 C. Bond, A. N. Santiago-Ruiz, Q. Tang, et al., “Technological advances in super-resolution microscopy to study cellular processes,” Molecular Cell 82(2), 315–332 (2022).

41 R. >D’Antuono, “Airyscan and confocal line pattern,” (2022).

42 D. Sage, T. A. Pham, H. Babcock, et al., “Super-resolution fight club: assessment of 2d and 3d single-molecule localization microscopy software,” Nature Methods 2019 16:5 16, 387–395 (2019).

43 S. Culley, D. Albrecht, C. Jacobs, et al., “Quantitative mapping and minimization of superresolution optical imaging artifacts,” Nature methods 15(4), 263–266 (2018).

44 T. Azuma and T. Kei, “Super-resolution spinning-disk confocal microscopy using optical photon reassignment,” Opt. Express 23, 15003–15011 (2015).

45 N. T. Feric, I. Pallotta, R. Singh, et al., “Engineered cardiac tissues generated in the biowire ii: A platform for human-based drug discovery,” Toxicological Sciences 172, 89–97 (2019).

46 W. J. de Lange, E. T. Farrell, C. R. Kreitzer, et al., “Human ipsc-engineered cardiac tissue platform faithfully models important cardiac physiology,” American Journal of Physiology-Heart and Circulatory Physiology 320(4), H1670–H1686 (2021).

47 Z. Wang, A. Bovik, H. Sheikh, et al., “Image quality assessment: from error visibility to structural similarity,” IEEE Transactions on Image Processing 13(4), 600–612 (2004).

48 A. Badon, S. Bensussen, H. J. Gritton, et al., “Video-rate large-scale imaging with multi-z confocal microscopy,” Optica 6(4), 389–395 (2019).

49 T. D. Weber, M. V. Moya, J. Mertz, et al., “High-speed, multi-z confocal microscopy for voltage imaging in densely labeled neuronal populations,” bioRxiv (2021).

50 B. Zhao, M. Koyama, and J. Mertz, “High-resolution multi-z confocal microscopy with a diffractive optical element,” Biomed. Opt. Express 14, 3057–3071 (2023).

51 H. Ma, F. Long, S. Zeng, et al., “Fast and precise algorithm based on maximum radial symmetry for single molecule localization,” Opt. Lett. 37, 2481–2483 (2012).

52 A. V. Kashchuk, O. Perederiy, C. Caldini, et al., “Particle localization using local gradients and its application to nanometer stabilization of a microscope,” ACS Nano 17(2), 1344–1354 (2023).

53 I. J. Schoenberg, Cardinal spline interpolation, SIAM (1973).

54 N. Kanopoulos, N. Vasanthavada, and R. L. Baker, “Design of an image edge detection filter using the sobel operator,” IEEE Journal of solid-state circuits 23(2), 358–367 (1988).

55 J. Pawley, Handbook of biological confocal microscopy, vol. 236, Springer Science & Business Media (2006).

56 C. N. Toepfer, A. C. Garfinkel, G. Venturini, et al., “Myosin sequestration regulates sarcomere function, cardiomyocyte energetics, and metabolism, informing the pathogenesis of hypertrophic cardiomyopathy,” Circulation 141(10), 828–842 (2020).

57 X. Lian, J. Zhang, S. M. Azarin, et al., “Directed cardiomyocyte differentiation from human pluripotent stem cells by modulating wnt/*β*-catenin signaling under fully defined conditions,” Nature protocols 8(1), 162–175 (2013).

58 J. Javor, S. Sundaram, C. Chen, et al., “Controlled strain of cardiac microtissue via magnetic actuation,” in 2020 IEEE 33rd International Conference on Micro Electro Mechanical Systems (MEMS), 452–455, IEEE (2020).

