## Supplementary Material for "Resolution enhancement with deblurring by pixel reassignment (DPR)"

**Bingying Zhao**[a,*], **Jerome Mertz**[b]
[a]Department of Electrical and Computer Engineering, Boston University, MA 02215,
[b]Department of Biomedical Engineering, Boston University, MA 02215

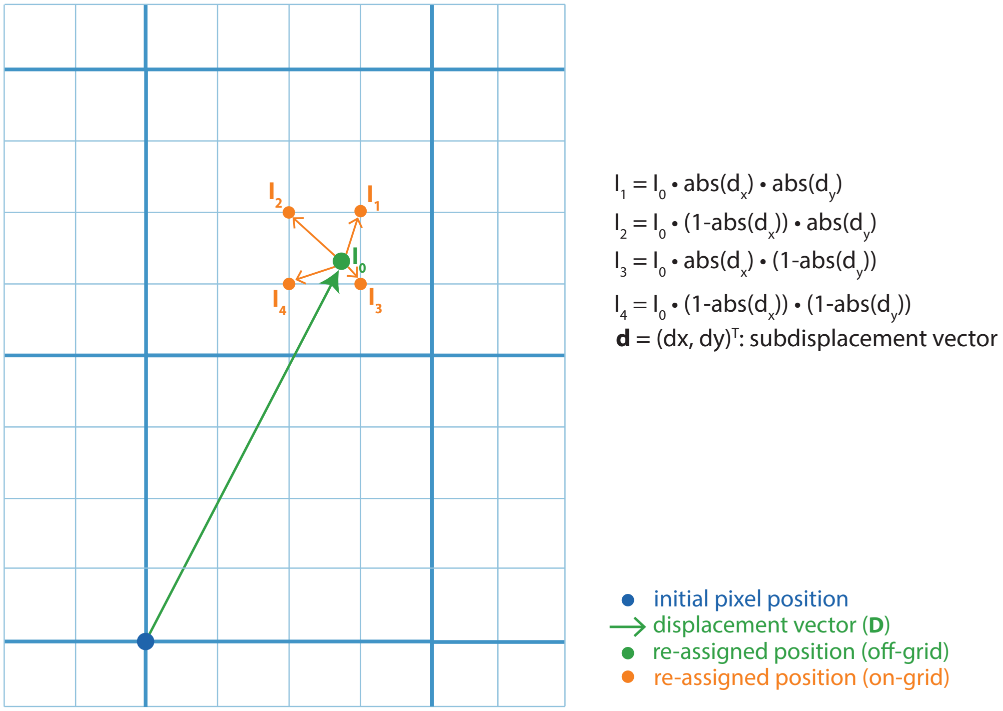


**Fig S1** The working principle of weighted pixel reassignment.

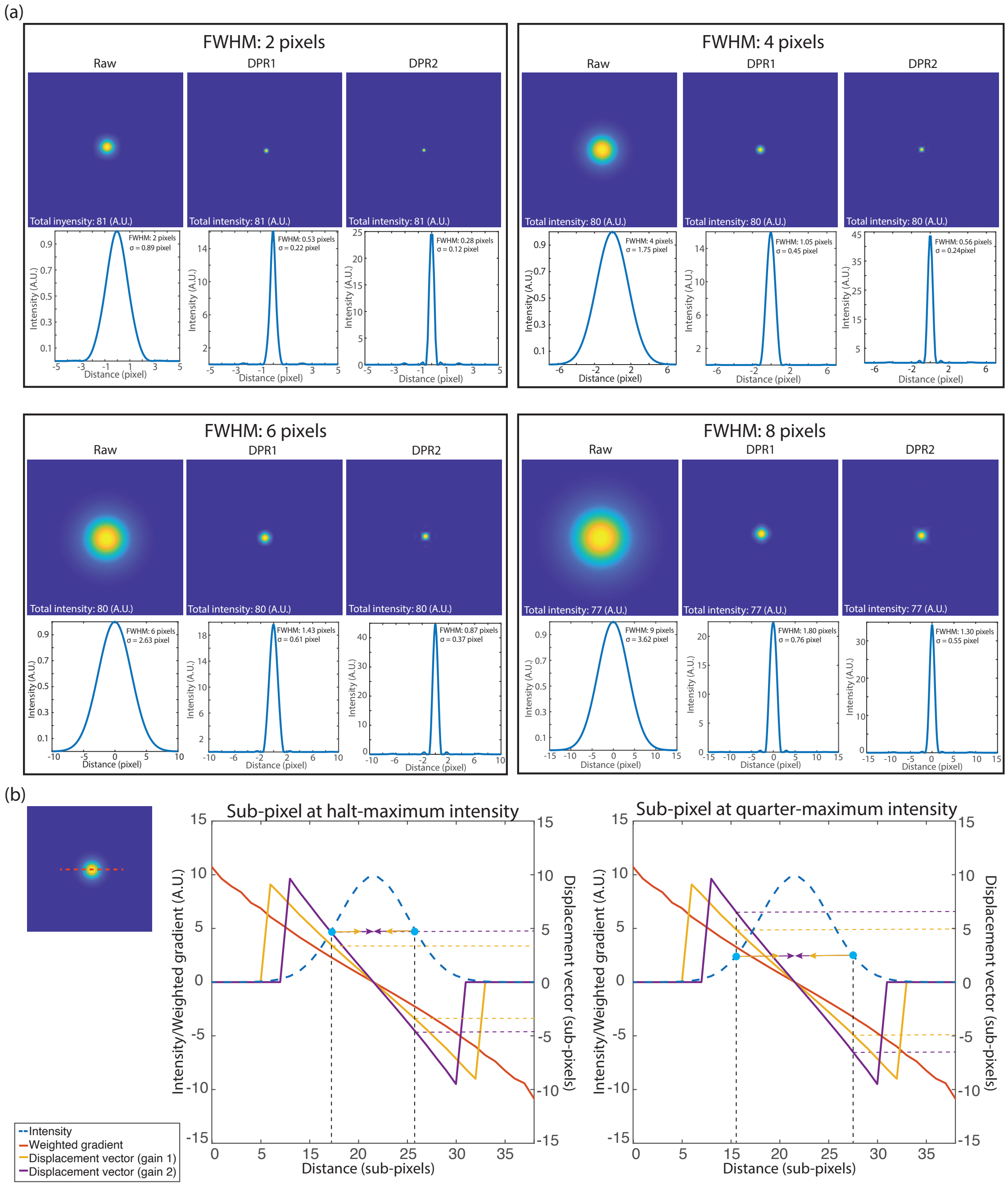


**Fig S2** DPR applied to simulated Gaussian PSFs of different sizes with gains 1 and 2. (a) The intensity images and horizontal line profiles of PSFs (with FWHMs of 2, 4, 6, and 8 pixels) before and after DPR gains 1 and 2. Total intensity: the sum of the intensity values at each sub-pixel in the image ('pixel' refers to raw-image pixel size). FWHM on the line profile: calculated FWHM after Gaussian fitting. $\sigma$ on the line profile: calculated RMS with Gaussian fitting. PSF FWHM: same as used in simulation, local-minimum filter radius: 4 times the FWHM. (b) The correlation between displacement vectors and different gains. Left: raw intensity image of a PSF whose FWHM is 4 pixels. Middle and right: the profiles for raw intensity, weighted gradient, and displacement vector under gain 1 and 2 along the red dashed line indicated on the left; yellow arrows indicate the displacement vectors for sub-pixels located at the half-maximum intensity (middle) and at the quarter-maximum intensity (right) with DPR gain 1; purple arrows indicate the displacement vectors for sub-pixels located at the half-maximum intensity (middle) and at the quarter-maximum intensity (right) with DPR gain 2.

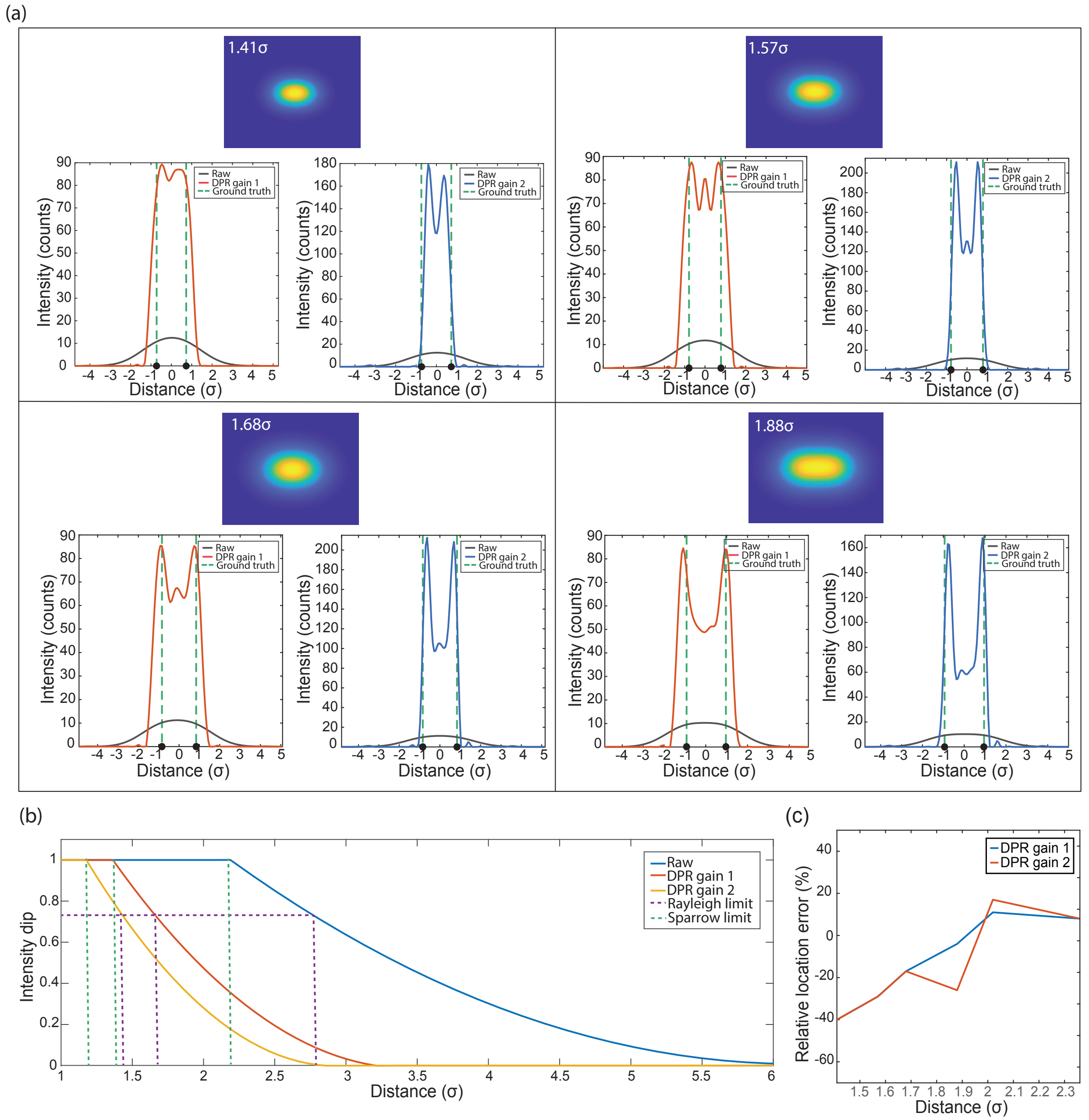


**Fig S3** The separation accuracy of DPR. (a) DPR applied to simulated images of two point objects separated by different distances. PSF FWHM: 2.35$\sigma$, local-minimum filter radius: 8$\sigma$. (b) The resolution enhancements of DPR gain 1 and DPR gain 2. (c) The separation errors resulting from DPR.

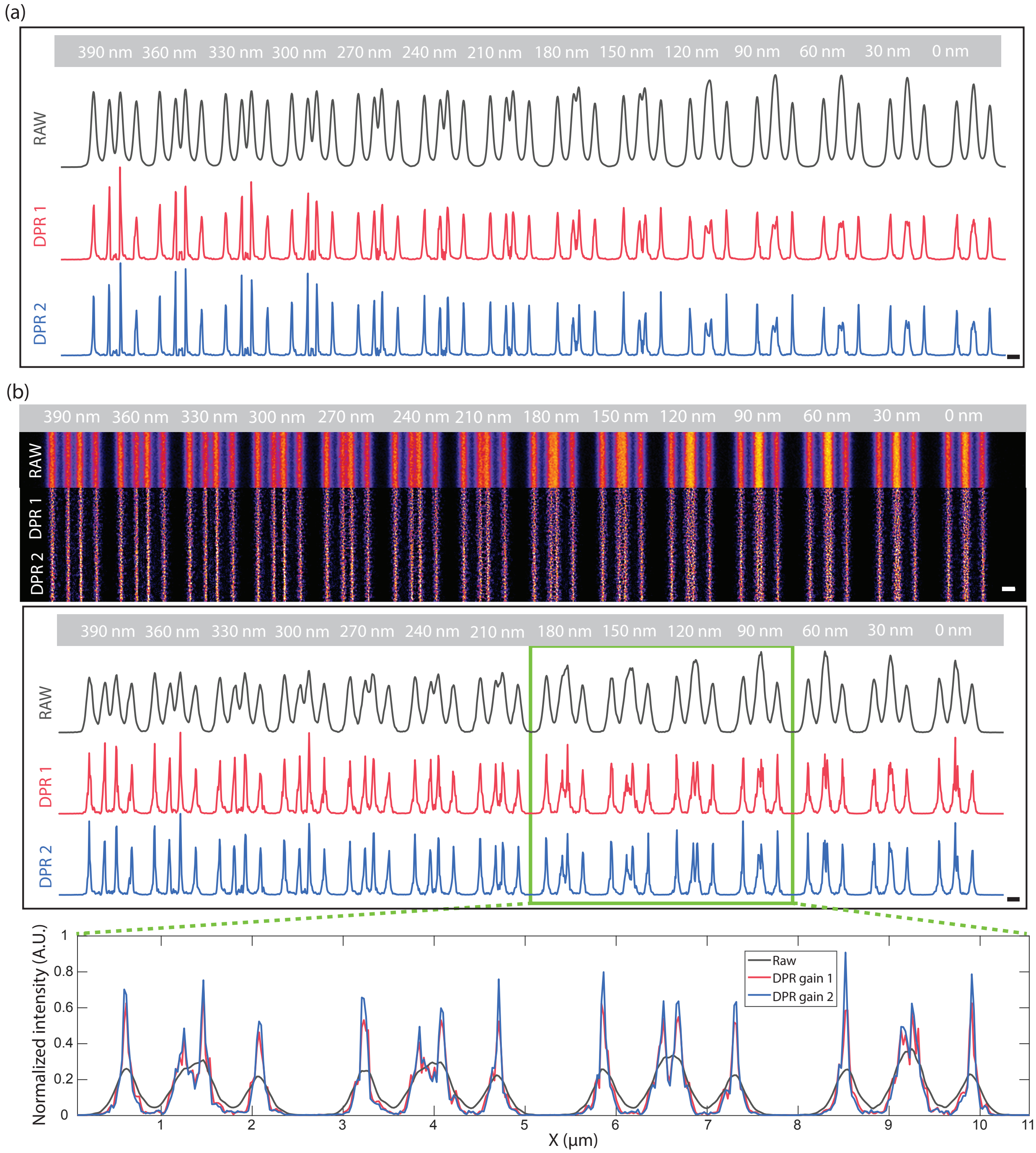


**Fig S4** DPR applied to fluorescent lines of increasing separation from 0 nm to 390 nm. (a) Raw data acquired by an Airyscan microscope. The full set of line intensity profiles for raw, DPR gain 1, and DPR gain 2. Scale bar: 500 nm. PSF FWHM: 4 pixels, local-minimum filter radius: 7 pixels. (b) Raw data acquired by a confocal microscope. Top: the full set of line images for raw, DPR gain 1, and DPR gain 2. Middle: the full set of line intensity profiles for raw, DPR gain 1, and DPR gain 2. Bottom: the expanded view of four line profiles for raw, DPR gain 1, and DPR gain 2, indicated by the green rectangle. PSF FWHM: 5 pixels, local-minimum filter radius: 9 pixels. Scale bar: 500 nm.

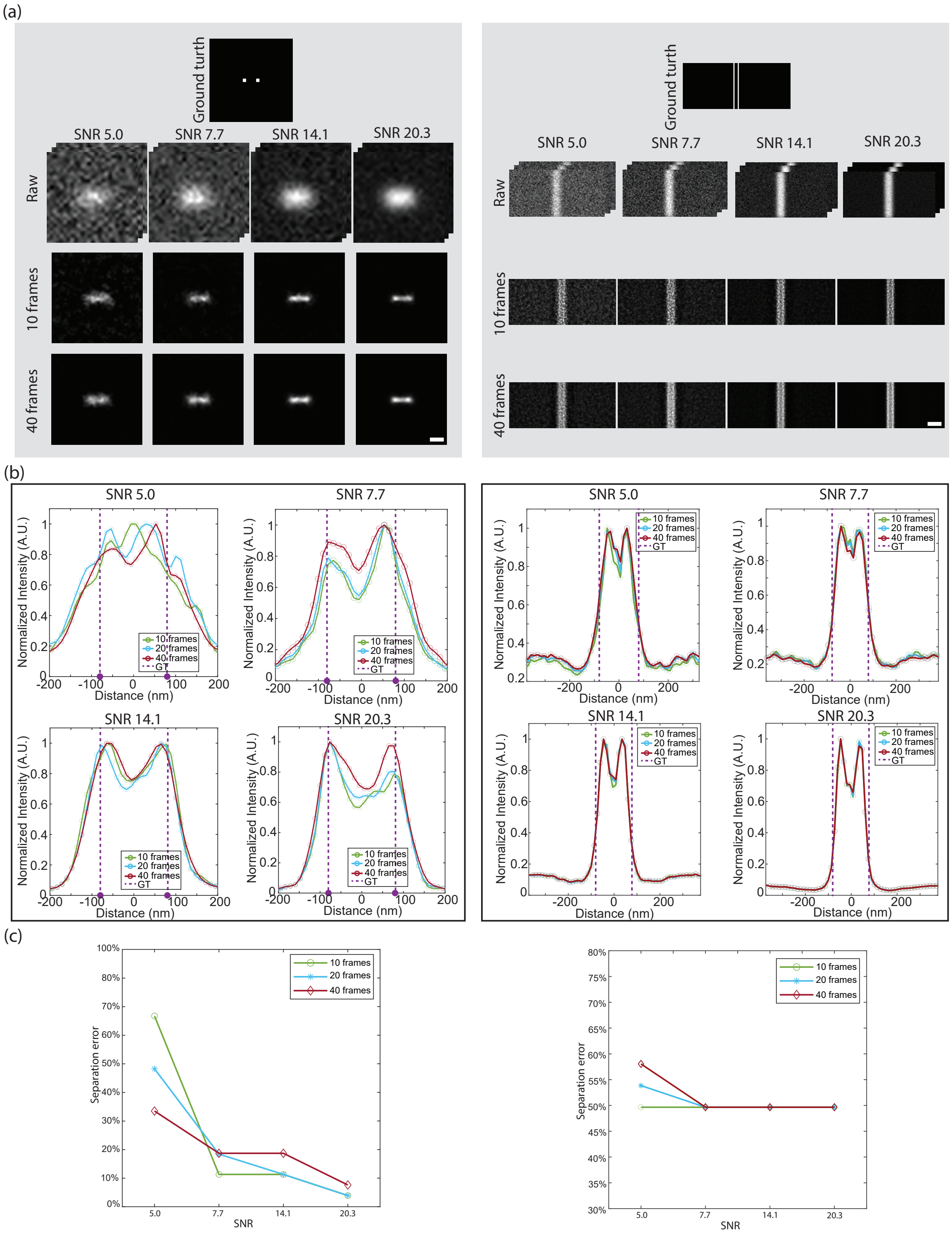


**Fig S5** DPR applied to simulated images of different SNRs. (a) Intensity images with different SNRs of the raw simulated two point objects (left) separated by 160 nm and two line objects (right) separated by 160 nm, without and with DPR. Scale bar: 100 nm. PSF FWHM: 5 pixels, local-minimum filter radius: 17 pixels, gain: 1. (b) Intensity profiles and (c) separation errors of the DPR reconstructed images when averaging over different numbers of frames. Dashed purple lines represent the ground-truth positions. Left: two point objects. Right: two line objects.

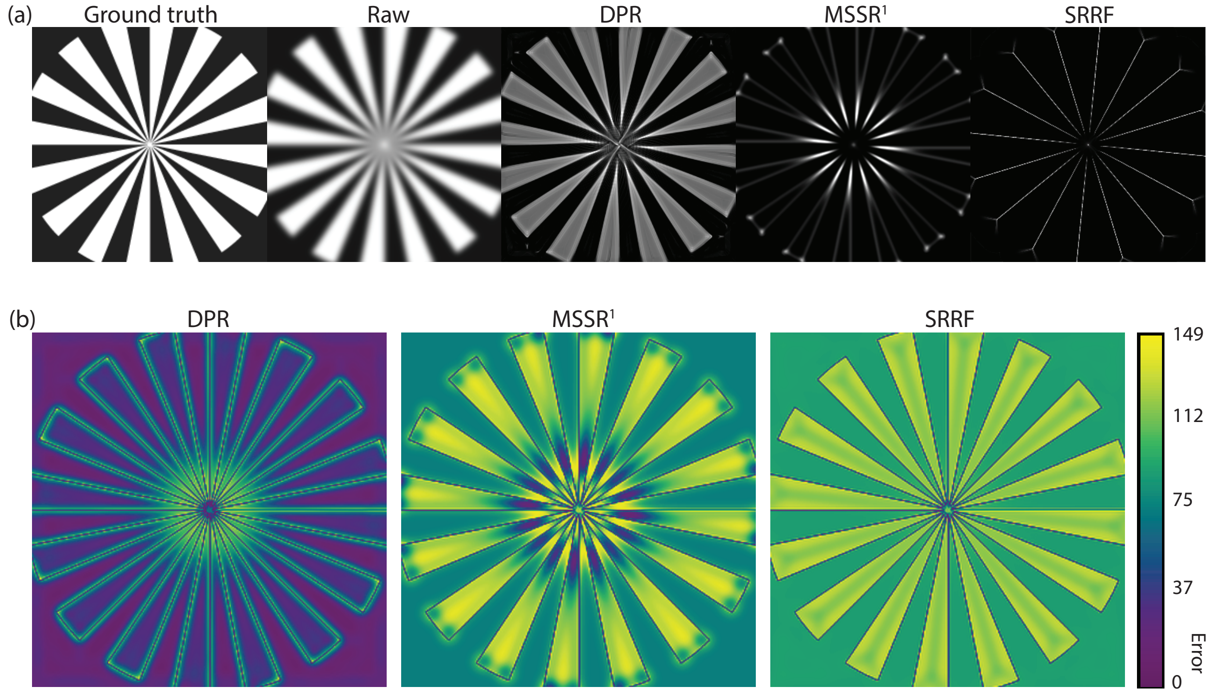


**Fig S6** Simulated Siemens star target. (a) The ground truth, simulated raw image, DPR-enhanced images, the first order of MSSR ($MSSR^1$)-enhanced image, and SRRF-enhanced image. DPR parameters: PSF FWHM 6 pixels, local-minimum filter radius: 13 pixels, gain 1. MSSR parameters: PSF FWHM 6 pixels, magnification 2. SRRF parameters: ring radius 0.5, magnification 2, axes 6. (b) The resolution scaled error maps for DPR-enhanced images, $MSSR^1$-enhanced image, and SRRF-enhanced image. NanoJ-SQUIRREL parameters: Ground truth selected as the reference image; representative resolution scaling function (RSF) set as unknown; Max Mag. in Optimization set as 5.

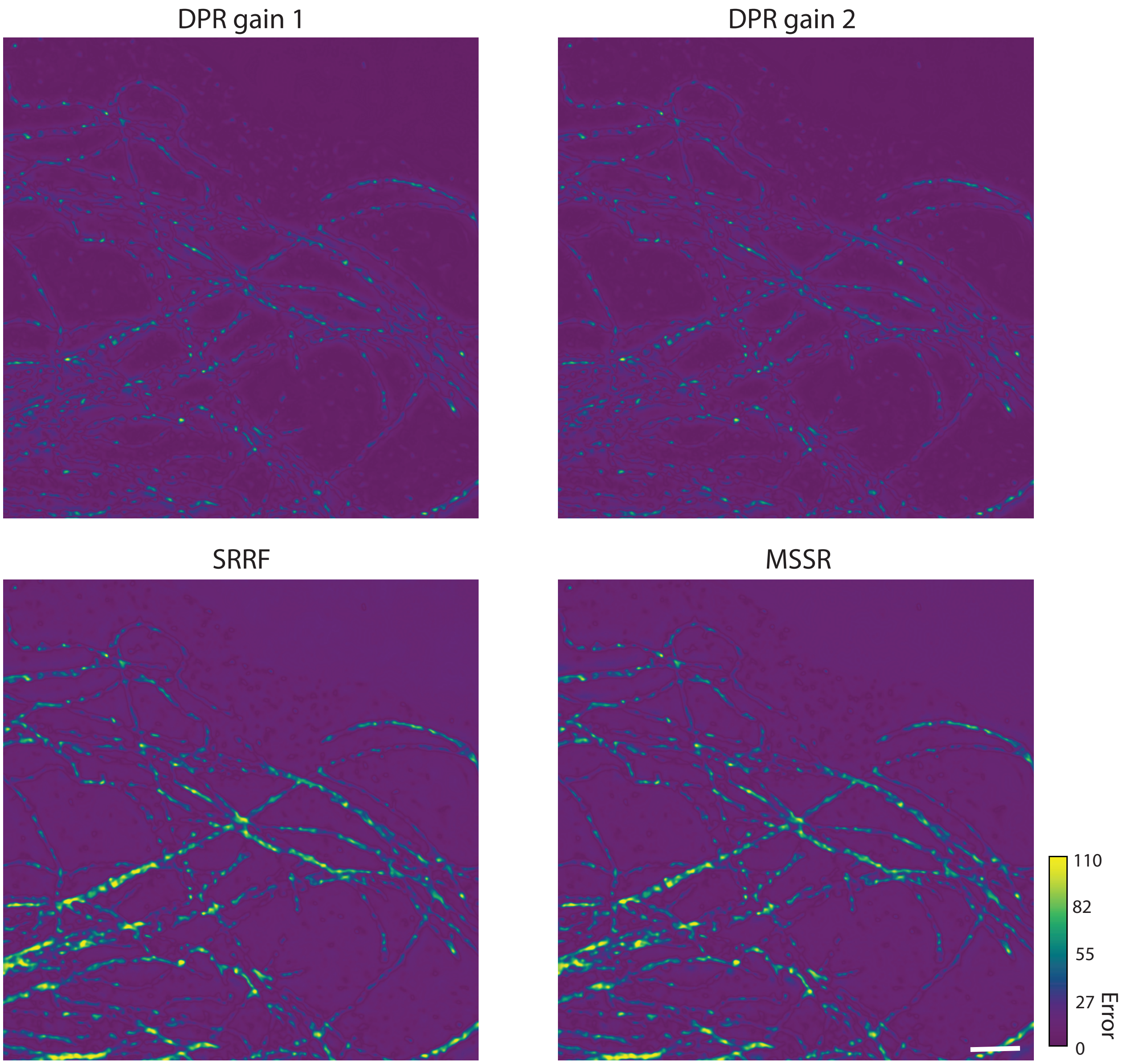


**Fig S7** The resolution scaled error maps for Fig. 5. NanoJ-SQUIRREL parameters: Ground truth selected as the SoRa image; representative RSF set as unknown; Max Mag. in Optimization set as 5. Scale bar: 2 $\mu$m.

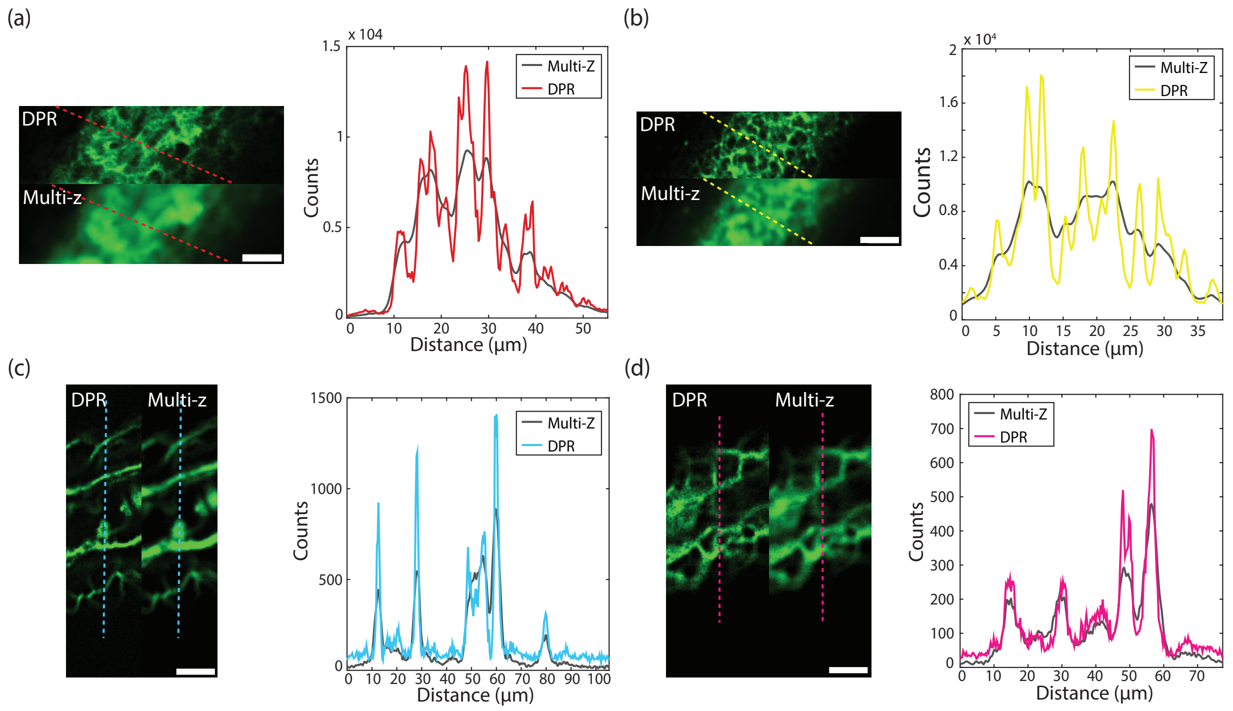


**Fig S8** Zoomed-in ROIs of multi-z confocal zebrafish imaging. (a) ROI indicated by the red rectangle in Fig. 7(a), and intensity profile along the red dashed line. Scale bar: 10 $\mu$m. (b) ROI indicated by the yellow rectangle in Fig. 7(a), and intensity profile along the yellow. Scale bar: 12 $\mu$m. (c) ROI indicated by the cyan rectangle in Fig. 7(b), and intensity profile along the cyan dashed line. Scale bar: 18 $\mu$m. (d) ROI indicated by the magenta rectangle in Fig. 7(b), and intensity profile along the magenta dashed line. Scale bar: 13 $\mu$m

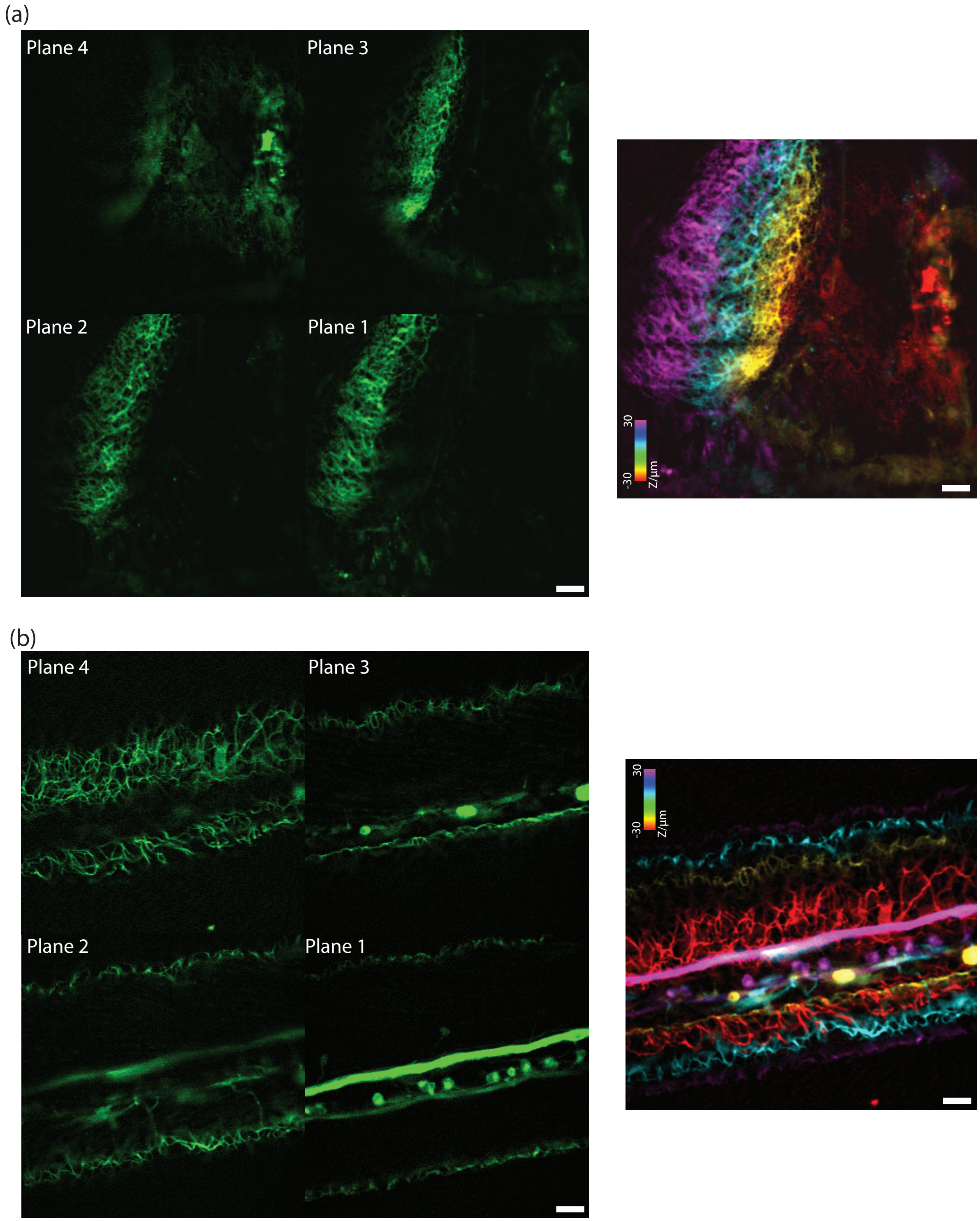


**Fig S9** Comparison high-resolution multi-z confocal zebrafish imaging, as reproduced from Zhao et. al., Biomed. Opt. Express 14, 3057 (2023). (a) *In-vivo* multiplane images of the brain region in a zebrafish 6 dpf. Left: four image planes. Right: merged with color corresponding to depth. (b) In-vivo multiplane images of the zebrafish tail region in the same zebrafish. Left: four image planes. Right: merged with color corresponding to depth. Plane 1-4: the deepest to the shallowest. Inter-plane separation: 20 $\mu$m. Scale bar: 20 $\mu$m

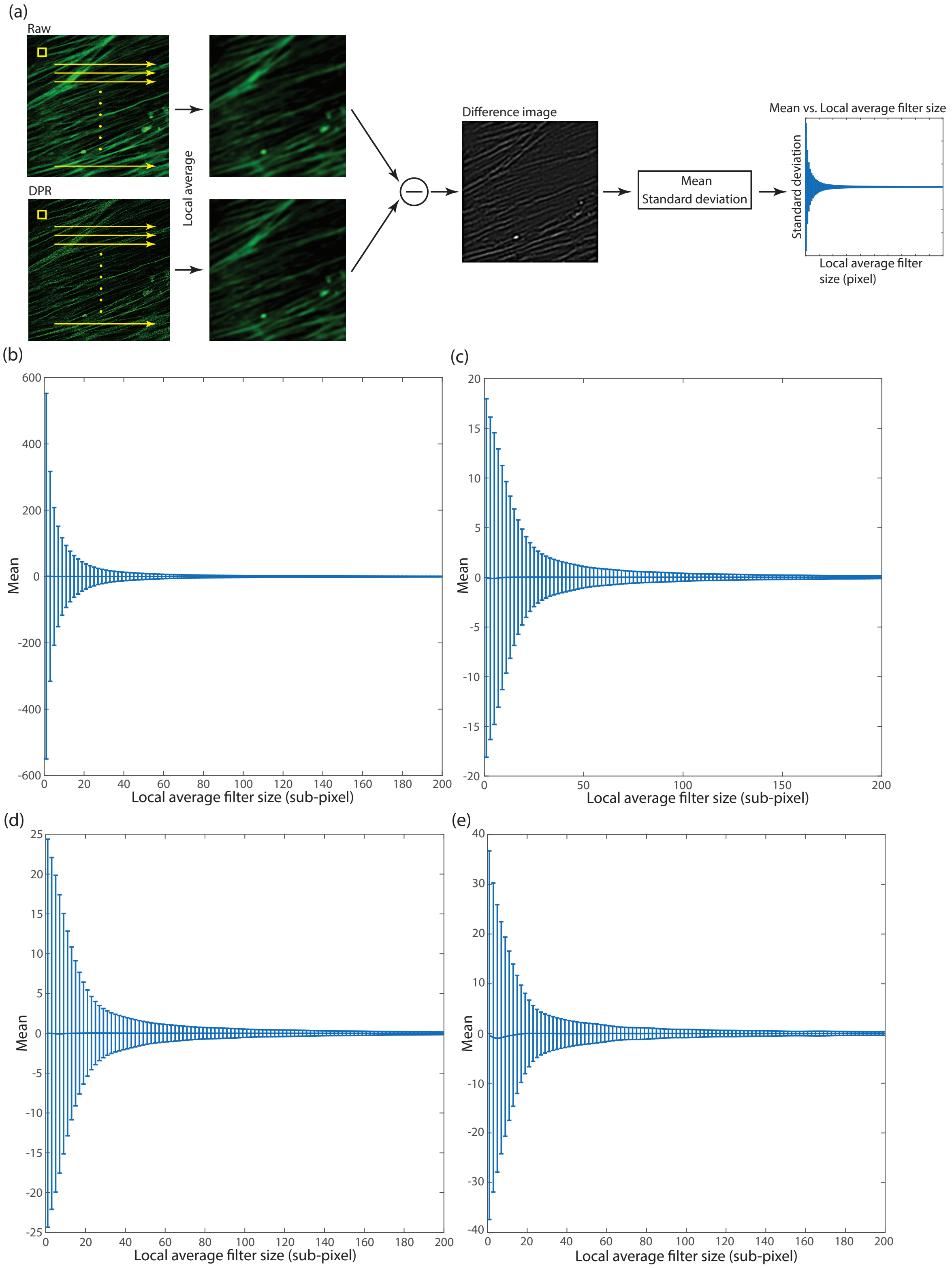


**Fig S10** Evaluation of preservation of local intensity with DPR (a) Evaluation procedure. Mean values of the difference images for Fig. 4 - confocal images of BPAE cells after DPR (b), Fig. 5 - SoRa images of BPAE cells after DPR gain 1 (c), Fig. 5 - SoRa images of BPAE cells after DPR gain 2 (d), and Fig. 6(a) - simulated images of sarcomere after DPR (e), with standard deviation shown as error bars.

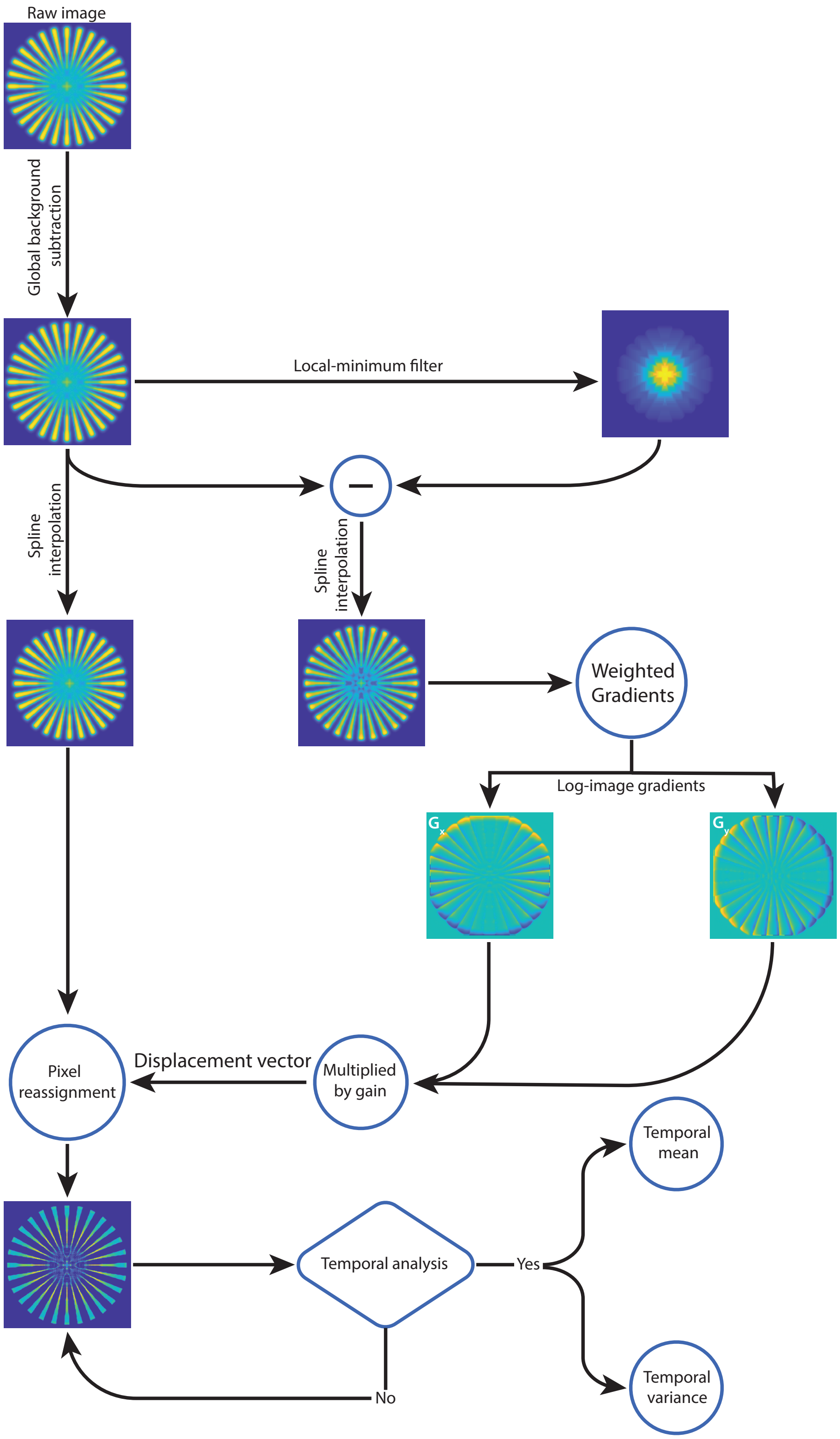


**Fig S11** The overall workflow of DPR.

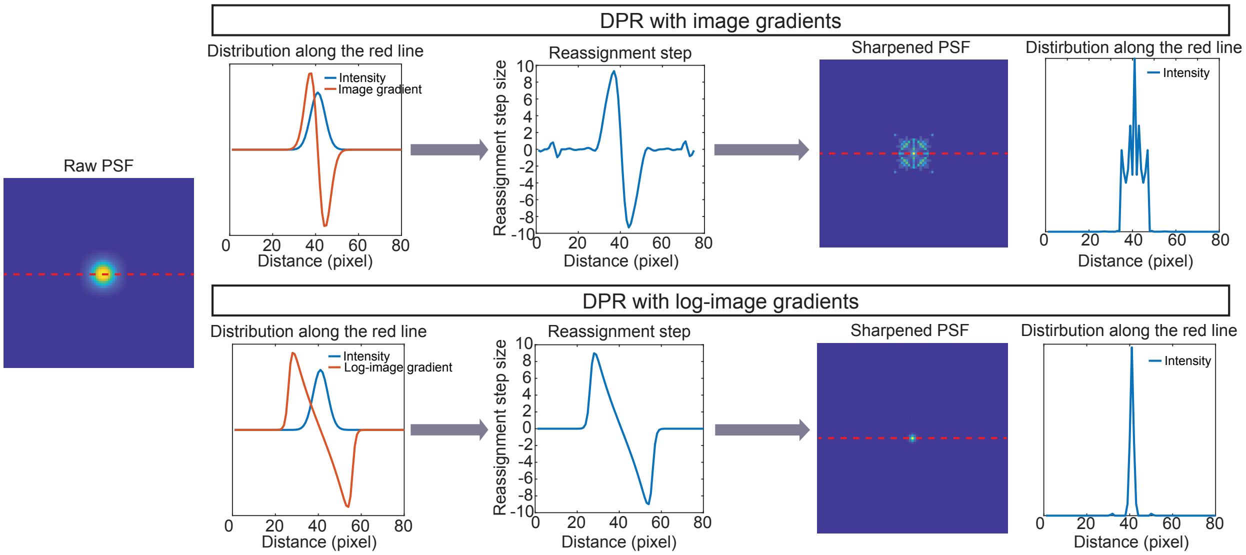


**Fig S12** DPR using gradients compared to with log-PSF gradients.

**Table S1** Total counts of images (intensities)

| Figure | Description | Raw | DPR gain 1 | DPR gain 2 | MSSR | SRRF |
|---|---|---|---|---|---|---|
| Fig. 1(a) | Simulated single PSF | 801 | 801 | - | - | - |
| Fig. 2(a) | Simulated two point objects separated by 1.68$\sigma$ | $1.24\times10^4$ | $1.24\times10^4$ | $1.24\times10^4$ | - | - |
| | Simulated two point objects separated by 1.41$\sigma$ | $1.27\times10^4$ | $1.27\times10^4$ | $1.27\times10^4$ | - | - |
| Fig. 2(b) | Fluorescent lines | $3.39\times10^8$ | $3.39\times10^8$ | $3.39\times10^8$ | - | - |
| Fig. 3(c) | SMLM challenge 2016 (temporal mean) | $3.87\times10^6$ | $3.87\times10^6$ | $3.87\times10^6$ | - | - |
| Fig. 4(a) | Confocal images of BPAE cells | $5.37\times10^8$ | - | $5.36\times10^8$ | $2.39\times10^7$ | $3.99\times10^7$ |
| Fig. 5(a) | SoRa images of BPAE cells | $3.28\times10^7$ | $3.28\times10^7$ | $3.27\times10^7$ | $2.38\times10^6$ | $2.98\times10^6$ |
| Fig. 6(a) | Simulated hiPSC-CMs images (temporal mean) | $2.54\times10^7$ | $2.54\times10^7$ | - | - | - |
| Fig. 6(b) | Low-resolution images of hiPSC-CMT organoids (temporal mean) | $4.63\times10^7$ | $4.62\times10^7$ | - | - | - |
| Fig. 6(c) | High-resolution images of hiPSC-CMT organoids (temporal mean) | $1.88\times10^8$ | $1.87\times10^8$ | - | - | - |
| Fig. 7(a) | Plane 1 | $4.01\times10^8$ | $4.00\times10^8$ | - | - | - |
| | Plane 2 | $3.36\times10^8$ | $3.36\times10^8$ | - | - | - |
| | Plane 3 | $1.91\times10^8$ | $1.91\times10^8$ | - | - | - |
| | Plane 4 | $1.29\times10^8$ | $1.29\times10^8$ | - | - | - |
| Fig. 7(b) | Plane 1 | $1.30\times10^8$ | $1.30\times10^8$ | - | - | - |
| | Plane 2 | $8.40\times10^7$ | $8.37\times10^7$ | - | - | - |
| | Plane 3 | $8.17\times10^7$ | $8.14\times10^7$ | - | - | - |
| | Plane 4 | $4.36\times10^7$ | $4.34\times10^7$ | - | - | - |

**Table S2** Parameters used in DPR, MSSR, and SRRF in the Results

| Figure | DPR | | | | MSSR | | | SRRF | | |
|---|---|---|---|---|---|---|---|---|---|---|
| | PSF FWHM | Local-minimum filter radius | Gain | Temporal analysis | PSF FWHM | Magnification | Order | Ring radius | Magnification | Axes |
| Fig. 2(a) | $2.35\sigma$ | $8\sigma$ | 1&2 | None | - | | | | | |
| Fig. 2(b) | 4 pixels | 7 pixels | 1&2 | None | - | | | | | |
| Fig. 3 | 2.7 pixels | 5 pixels | 1&2 | Temporal average & Temporal variance | - | | | | | |
| Fig. 4 | 4 pixels | 12 pixels | 2 | None | 4 pixels | 2 | 1 | 0.5 | 4 | 6 |
| Fig. 5 | 2 pixels | 40 pixels | 1&2 | None | 2 pixels | 4 | 1 | 0.5 | 4 | 6 |
| Fig. 6(a) | 4 pixels | 7 pixels | 1 | None | - | | | | | |
| Fig. 6(b) | 4 pixels | 7 pixels | 1 | Temporal average | - | | | | | |
| Fig. 6(c) | 2 pixels | 4 pixels | 1 | Temporal average | - | | | | | |
| Fig. 7 | 5 pixels | 25 pixels | 1 | Temporal average | - | | | | | |

**Algorithm S1** DPR

1: **Input:** $I$ = raw image, $\delta$ = PSF FWHM, $g$ = gain, $r$ = local-minimum filter radius, $T$ = temporal process
2: **Initialization:** $f_{local} \leftarrow$ local-minimum filter, $g_x, g_y \leftarrow$ Sobel kernel, $Gaussian(10) \leftarrow$ Gaussian function with RMS of 10
3: **Readout image size:** $n_x, n_y, n_f \leftarrow$ size of $I$
4: **Upscale factor:** $M_x, M_y \leftarrow \frac{\delta}{8} \times n_x, \frac{\delta}{8} \times n_y$
5: **for** $i$ = 1, 2,...,$n_f$ **do**
6: $\quad I_{in} \leftarrow i$th frame of $I$
7: $\quad$ **Global background subtraction:** $I_{in} \leftarrow I_{in} - min(I_{in})$
8: $\quad$ **Local minimum:** $L_{min} \leftarrow I_{in} \otimes f_{local}$
9: $\quad$ **Local minimum subtraction:** $I_{eq} \leftarrow I_{in} - L_{min}$
10: $\quad$ **Interpolation:** $I_{mag:in}, I_{mag:eq} \leftarrow$ Upscale $I_{in}$ and $I_{eq}$ by $M_x$ times horizontally and $M_y$ times vertically
11: $\quad$ **Local normalization:** $I_{mag:eq} \leftarrow I_{mag:eq}/(I_{mag:eq} \otimes Gaussian(10) + 10^{-5})$
12: $\quad$ **Gradients:** $G_x, G_y \leftarrow I_{mag:eq} \otimes g_x, I_{mag:eq} \otimes g_y$
13: $\quad$ **Weighted gradients:** $G_x, G_y \leftarrow \frac{G_x}{I_{mag:eq}}, \frac{G_y}{I_{mag:eq}}$
14: $\quad$ **Reassignment step sizes:** $D_x, D_y \leftarrow G_x \times g, G_y \times g$
15: $\quad$ **Weighted pixel reassignment** on $I_{mag:in}$
16: **end for**
17: **if** T=None **then**
18: $\quad$ Output reconstructed image stack
19: **else if** T=Average **then**
20: $\quad$ Output the temporal average of the reconstructed images
21: **else if** T=variance **then**
22: $\quad$ Output the temporal variance of the reconstructed images
23: **end if**

2: Sobel filters

$$g_x = \begin{bmatrix} +1 & 0 & -1 \\ +2 & 0 & -2 \\ +1 & 0 & -1 \end{bmatrix} \tag{S1}$$

$$g_y = \begin{bmatrix} +1 & +2 & +1 \\ 0 & 0 & 0 \\ +1 & -2 & -1 \end{bmatrix} \tag{S2}$$

10: Interpolation function

$$I_{mag:in}(x',\, y',\, n) = spline(I_{in}(x,\, y,\, n)) \tag{S3}$$

$$I_{mag:eq}(x',\, y',\, n) = spline(I_{eq}(x,\, y,\, n)) \tag{S4}$$

14: Reassignment step sizes

$$D_x(x',\, y'\, n) = \begin{cases} G_x(x',\, y',\, n) \cdot g, & \text{if } G_x(x',\, y',\, n) \cdot g \leq 10 \\ 0, & \text{if } G_x(x',\, y',\, n) \cdot g > 10 \end{cases} \tag{S5}$$

$$D_y(x',\, y',\, n) = \begin{cases} G_y(x',\, y',\, n) \cdot g, & \text{if } G_y(x',\, y',\, n) \cdot g \leq 10 \\ 0, & \text{if } G_y(x',\, y',\, n) \cdot g > 10 \end{cases} \tag{S6}$$

15: Method of weighted pixel reassignment to the nearest pixels

Sub-displacement vector $\mathbf{d} = (d_x\mathbf{x},\, d_y\mathbf{y})$:

$$d_x = D_x - fix(D_x) \tag{S7}$$

$$d_y = D_y - fix(D_y) \tag{S8}$$

The weighted distribution of a pixel with intensity $I_0$ located at ($x'_0$, $y'_0$) is given by

$$\begin{aligned} I_{mag:in}(ceil(x'_0 + D_x), ceil(y'_0 + D_y)) =& I_{mag:in}(ceil(x'_0 + D_x), ceil(y'_0 + D_y)) \\ &+ I_0 \cdot abs(d_x)abs(d_y) \end{aligned} \tag{S9}$$

$$\begin{aligned} I_{mag:in}(floor(x'_0 + D_x), ceil(y'_0 + D_y)) =& I_{mag:in}(floor(x'_0 + D_x), ceil(y'_0 + D_y)) \\ &+ I_0 \cdot (1 - abs(d_x))abs(dy) \end{aligned} \tag{S10}$$

$$\begin{aligned} I_{mag:in}(ceil(x'_0 + D_x), floor(y'_0 + D_y)) =& I_{mag:in}(ceil(x'_0 + D_x), floor(y'_0 + D_y)) \\ &+ I_0 \cdot abs(d_x)(1 - abs(d_y)) \end{aligned} \tag{S11}$$

$$\begin{aligned} I_{mag:in}(floor(x'_0 + D_x), floor(y'_0 + D_y)) =& I_{mag:in}(floor(x'_0 + D_x), floor(y'_0 + D_y)) \\ &+ I_0 \cdot (1 - abs(d_x))(1 - abs(d_y)) \end{aligned} \tag{S12}$$
